# Integrated multi-omics profiling of gut bacterial extracellular vesicles links cargo composition to host transcriptional responses

**DOI:** 10.64898/2026.07.01.732738

**Authors:** Adarsh Singh, Yanjia Zhang, Jiucheng Ding, Qiaojuan Shi, Anas Saleh, Josh Jones, Xieyue Xiao, Yi-Yuan Lee, Kevin B. Weyant, Matthew P. DeLisa, Aaron Timperman, Kyu Y. Rhee, Ilana L. Brito

**Affiliations:** Nancy E. and Peter C. Meinig School of Biomedical Engineering, Cornell University, Ithaca, NY, USA; Department of Bioengineering, University of Pennsylvania, Philadelphia, PA, USA; Biochemistry and Biophysics, University of Pennsylvania, Philadelphia, PA, USA; Department of Medicine, Weill Cornell Medicine, New York, NY, USA; Robert F. Smith School of Chemical and Biomolecular Engineering, Cornell University, Ithaca, NY, USA; Cornell Institute of Biotechnology, Cornell University, Ithaca, NY, USA

## Abstract

Bacterial extracellular vesicles (bEVs) enable gut microbiota to deliver bioactive cargo to host cells, yet the specific bacterial producers have not been systematically identified. Here, we profiled stool-derived bEVs from healthy individuals using metaproteomic profiling, revealing that vesiculation is widespread across gut bacterial phyla. We selected a subset of vesiculating species, showing that bEVs localize to distinct tissues *in vivo*, including extraintestinal sites such as lung, liver, kidney, and bone, suggesting roles beyond the gastrointestinal tract. We profiled intestinal epithelial cells and macrophages after endocytosing bEVs from various species, uncovering pronounced, cell-type-specific responses. For example, *Bacteroides fragilis* bEVs promote anti-inflammatory mitochondrial-telomeric regulation, while a set of commensal-derived bEVs contribute to epithelial survival and structural renewal. By combining proteomic, lipidomic, and metabolomic cargo profiling with transcriptional output, we find that specific Bacteroidota-derived protein cargo activates cytoprotective stress defenses while attenuating inflammatory signaling. Together, these findings establish a multi-layered comparative atlas of bEV composition, uptake, and host response, providing a framework for understanding bEV-mediated microbiome-host communication.

## Introduction

The gut microbiome can directly affect host tissues, with one mechanism being bacterial extracellular vesicles (bEVs). bEVs enable bacteria to influence intestinal epithelial cells, as well as extraintestinal tissues, such as the liver^1^ and joints^2^, without direct contact. However, thus far, studies on these nanoscale, membrane-bound particles have been limited to single or a few bacterial species.

During vesiculation, bEVs encapsulate a defined subset of bacterial components, including proteins, lipids, small molecules, and nucleic acids that are dependent on their structures. Gram-negative bacteria are capable of producing outer membrane vesicles (OMVs) and outer-inner membrane vesicles (OIMVs), while Gram-positive bacteria release only inner/cytoplasmic membrane vesicles (IMVs/CMVs)^3^. bEV cargo can be functionally specialized, encompassing factors involved in immune modulation, oxidative stress resistance, and virulence^4–6^.

bEVs can engage host cells either through surface interactions, where they bind pattern-recognition receptors and trigger signaling without internalization, or through endocytic uptake that delivers bEV cargo into host cells^7–9^. After endocytosis, bEV cargo may escape the endosome or directly interact with host components through cytosolic delivery. For example, lipopolysaccharide (LPS) present on enterohemorrhagic *Escherichia coli* (EHEC)-derived bEVs has been shown *in vitro* to escape the endosome to induce pyroptosis^10^; while OmpA from *Acinetobacter baumannii* causes mitochondrial fragmentation, leading to cell death^11^. *In vivo,* bEVs may traverse the gut mucosal barrier to deliver microbial cargo to host epithelial and immune cells, triggering both protective and pathogenic responses. For example, bEVs derived from *Bacteroides fragilis* carry the immunomodulatory capsular polysaccharide A (PSA), which induces T helper cells and protects from experimental colitis^12^. In contrast, *Bacteroides thetaiotaomicron* delivers sulfatases via bEVs, which exacerbate colitis^13^. bEVs have also been shown to translocate to different organs, including the liver, kidney, and spleen^1^. Notably, bEVs derived from the gut pathobiont *Fusobacterium nucleatum* localize to mouse paws, where they aggravate rheumatoid arthritis^2,14^.

Collectively, these studies underscore bEVs as important conduits for host-microbiome communication. Whereas the majority of existing work has focused on bEVs derived from pathogens, those derived from gut pathobionts and commensals are comparatively understudied. Systematically defining which members of the gut microbiome produce these bEVs is challenging. Estimates of bEV abundance in stool samples using 16S rDNA-based approaches are limited to family-level taxonomic resolution^15,16^ and may have low sensitivity due to the requirement for adequate representation of 16S rDNA in bEVs. Furthermore, comparisons of bEV cargo composition may vary substantially across studies, due to different isolation methods. Similarly, their reported pro-inflammatory or protective effects vary by experimental context^13,17–19^. As a result, the field lacks a systematic, species-resolved, multi-omic framework that functionally links bEV composition to host-cell uptake and functional responses.

Here, we systematically address the question of which bacteria produce bEVs in the human gut microbiome. We find that bEV-producing bacteria span diverse taxonomic groups. Based on these observations, we selected a representative set of commensal bacteria alongside known gut-associated pathobionts for targeted study, where we define their diversity, cargo composition, and functional impact on host cells. bEVs isolated from these bacteria exhibited marked differences in the mechanisms of uptake and caused pronounced differences in transcriptomic responses in bEV-treated intestinal epithelial cells and macrophages. Finally, integrative correlation analyses linked specific bEV features with host transcriptional responses, providing insight into how bEV composition shapes host-microbiome interactions.

## Methods

### Stool sample collection

A cohort of 11 people with no known chronic gastrointestinal disorders were recruited. All participants were consented under protocol #1911009200, approved by the Cornell University Institutional Review Board. Fresh stool samples were homogenized in equal weight 1:1 glycerol and PBS, and stored at -80⁰C until further processing.

### bEV isolation and purification from stool samples and bacterial culture

Frozen stool samples were homogenized, and 3 g of homogenate was resuspended in 10 mL phosphate-buffered saline (PBS). The suspension was centrifuged at 7,000 × g for 10 min at 4 °C, and the supernatant was transferred to a fresh tube. The remaining pellet was resuspended in an additional 10 mL PBS, and the centrifugation step was repeated for a total of three rounds. For bacterial cultures, strains were grown overnight in the appropriate growth medium, and cells were pelleted by centrifugation at 5,000 × g for 30 min at 4 °C. Supernatants from stool and bacterial cultures were filtered through 0.45 µm pore-size filters to remove residual cells and debris. Filtered supernatants were subjected to ultracentrifugation at 141,000 × g for 3 h at 4 °C to pellet bEVs, followed by washing with PBS. Crude bEV pellets were further purified using an iodixanol (OptiPrep™, Sigma) density gradient composed of 15%, 20%, 25%, 30%, 35%, 40%, and 45% (w/v) layers. Following ultracentrifugation, fractions 2-6 were collected and pooled to obtain purified bEVs, which were aliquoted and stored in PBS at −80 °C until further analysis. bEVs were inoculated in the respective media to check for live bacterial contamination. Protein concentration was determined using a bicinchoninic acid (BCA) assay according to the manufacturer’s instructions (Thermo Fisher Scientific). bEV size distribution and particle concentration were measured by nanoparticle tracking analysis (NTA) using a NanoSight NS300 system (Malvern) equipped with a 488 nm laser and sCMOS camera. Samples were appropriately diluted in PBS and introduced using a syringe pump set to 30 arbitrary units (AU). For each replicate, three 60 seconds videos were recorded and averaged for final analysis.

[utbl1]

### DNA extraction, shotgun metagenome sequencing, and taxonomy assessment

Total DNA was extracted from approximately 0.25 g of frozen stool using the DNeasy PowerSoil Kit (Qiagen) according to the manufacturer’s instructions. DNA concentration was determined with a Qubit fluorometer (Thermo Fisher Scientific), and 1 ng of input DNA was used for tagmentation and PCR amplification following the Nextera XT DNA Library Preparation protocol (Illumina). Libraries underwent double-size selection with AMPure XP beads (Beckman Coulter), and equimolar amounts of all libraries were pooled. The pooled library was sequenced on an Illumina NovaSeq platform using a paired-end 150-cycle run. Read Processing and Assembly Raw reads were dereplicated with PRINSEQ (v0.20.4) with -derep 12345, human-derived sequences were removed using BMTagger, and adapter sequences were trimmed with Trimmomatic ((v0.39) with ILLUMINACLIP:2:30:10:2:keepBothReads LEADING:3 TRAILING:3 MINLEN:36 to remove adapters ((ILLUMINACLIP:TruSeq3- PE.fa:2:30:10), remove leading low quality or N bases (below quality 3) (LEADING:3), remove trailing low quality or N bases (below quality 3) (TRAILING:3) and remove reads below 36 bases long (MINLEN:36)^20–22^. Processed reads from each sample were assembled using SPAdes in metagenomic mode (--meta) (v3.15.5) with k-mer sizes of 21, 29, 39, 59, 79 and 99^23,24^. The resulting contig was used an input for Kraken2 and Bracken for taxonomy assignment and beta-diversity abundance estimation.

### Stool bEV proteomics and taxonomy estimation

Stool bEV proteomics was performed at the Proteomics Resource Center (Rockefeller University). Briefly, samples were dried, and protein pellets were resuspended in 20 mM ammonium bicarbonate containing 1% NP-40. Proteins were precipitated overnight using cold acetone, and pellets were subsequently resuspended in 8 M urea for reduction and alkylation. Samples were digested sequentially with Lys-C followed by overnight trypsin digestion. Reactions were quenched with neat trifluoroacetic acid (TFA), and peptides were purified by solid-phase extraction prior to LC-MS analysis. Peptides were analyzed using a 70 min analytical gradient on a 12 cm C18 reversed-phase column-in-emitter configuration coupled to a Q Exactive HF mass spectrometer operated in high-resolution, high-mass-accuracy mode. Prior to analysis of individual samples, pooled samples were run to optimize acquisition parameters. To generate a reference database for protein identification, open reading frames (ORFs) were predicted from metagenomic contigs using Prodigal (v2.6.3)^25^. Predicted ORFs were functionally annotated by mapping to the UniRef90 database using DIAMOND^26^. Metagenome-derived ORFs were combined with the Unified Human Gastrointestinal Proteome (UHGP) database and clustered at 95% sequence identity using CD-HIT^27,28^. Representative centroid sequences were merged with the human UniProt-TrEMBL database to create the final search database^29^. Taxonomic assignment and protein identification were performed using the MetaLab2 workflow^30^. GO Biological Processes terms were extracted by mapping it to Uniref50, and similar terms were merged using REVIGO^31^. For the correlation analysis between metagenomic and proteomic datasets, discrepancies in taxonomic nomenclature arising from differences in database versions and naming conventions used by MetaPhlAn4 and MetaLab2 were first resolved. To ensure consistent taxonomic annotation across platforms, we developed a custom script to query the NCBI Taxonomy database and retrieve the current accepted NCBI taxonomic names corresponding to each identified lineage.

### TEM and SEM

For negative staining TEM of bEVs, formvar and carbon-coated 400 mesh copper grids were glow-discharged, loaded with 3-5 µL of sample for 1 minute, and excess liquid removed with filter paper. Grids were immediately stained with 1.5% aqueous uranyl acetate for 1 minute, air-dried for 5 minutes, and examined using a JEM-1400 TEM (JEOL USA, Peabody, MA) operated at 100 kV. Images were acquired using a Veleta 2K × 2K CCD camera (EM-SIS, Germany; WCM Microscopy and Image Analysis Core facility). For SEM of bacteria, 1 ml of overnight culture was pelleted and primary fixed with 2% Glutaraldehyde for 2 hours and further secondary fixed with 1% OsO4 for 1 hour. Samples underwent graded dehydration in ethanol (25%, 50%, 70%, 95%, 100%) and were transferred to Whatman filter paper for critical point drying using liquid CO₂ overnight. After drying, samples were sputter-coated with gold-palladium (60:40 ratio, ∼10 nm) for 60 seconds at 30 mA. Images were captured using a side-angle Everhart-Thornley secondary electron detector on a Zeiss Sigma 500 SEM (Cornell CCMR).

### bEV *in vivo* biodistribution

C57BL/6 mice (4–6 weeks old; Jackson Laboratory) were acclimatized for one week upon arrival, then transitioned to an alfalfa-free diet for an additional week before experimentation to minimize autofluorescence. bEVs were fluorescently labeled with Cy7 (Cytiva, 5 µM final concentration) per the manufacturer’s protocol, and free dye was removed by ultracentrifugation (141,000 × g, 20 min, TLA100.3 rotor) with three successive washes in 1xPBS. Mice were administered 25 µg (total protein) of Cy7-labeled bEVs or PBS (control) via oral gavage, and organs were harvested 12 hours post-gavage. *Ex vivo* fluorescence imaging was performed using an IVIS Spectrum imaging system (PerkinElmer), and signal intensity was quantified using Living Image software.

### Caco-2 endocytosis experiment

Caco-2 cells were seeded in 96-well plates (IBIDI) and maintained in DMEM supplemented with 10% fetal bovine serum (FBS) for 2-3 weeks to allow differentiation. On the day of the experiment, cells were pre-treated for 1 h under gentle shaking conditions with endocytosis inhibitors at the following final concentrations, as previously described : chlorpromazine-HCl (15 µg/mL, water), genistein (200 µM, DMSO), dynasore (80 µM, DMSO), methyl-β-cyclodextrin (10 mM, water), cytochalasin D (1 µg/mL, DMSO), and amiloride (0.1 mM, DMSO)^1,32^. For DMSO-solubilized inhibitors, the final DMSO concentration was diluted at least 1:100 to minimize solvent-related toxicity. Following inhibitor pre-treatment, cells were incubated with 0.5 µg DiD-labeled bEVs for 4 h^33^. Cells were detached using TrypLE Express (Thermo Fisher Scientific) and stained with Fixable Viability Dye (Invitrogen) to exclude dead cells. Cells were subsequently fixed with 4% paraformaldehyde (PFA). All treatment and staining steps were followed by three washes with 1× PBS, with 5 min incubation between washes. Samples were acquired on a Thermo Fisher Attune NxT flow cytometer at the Cornell Flow Cytometry Core Facility. During analysis, cells were sequentially gated to exclude debris, doublets, and non-viable cells. Gates were defined such that the percentage of DiD⁺ events in the no-bEV, no-drug control condition remained below 0.5%. Percent suppression of bEV uptake was calculated relative to the no-inhibitor control condition.

### Interaction study with Caco-2 and primary macrophage

Human primary monocytes were obtained from de-identified donors (University of Nebraska Medical Center) and differentiated into macrophages as previously described^34^. Briefly, monocytes were cultured in complete DMEM supplemented with human serum and differentiated for 7 days in the presence of 25 ng/mL M-CSF and 25 ng/mL GM-CSF, with media replacement every 2-3 days. Prior to experiments, cells were washed with 1× DPBS and maintained in antibiotic-free media. Caco-2 cells were cultured as described above. For bEV exposure experiments, both differentiated primary macrophages and Caco-2 cells were treated with 5 µg of purified bEvs. After 4 h of incubation, supernatants were removed, and cells were washed with PBS, lysed in TRIzol reagent, and stored at −80 °C until RNA extraction.

### RNA extraction and sequencing

We followed the TREx protocol provided by Cornell. Briefly, 200 µL of chloroform was added per 1 mL of Trizol, followed by vigorous shaking by hand for 15 seconds. The samples were centrifuged at 16,000 x g for 15 minutes at 4°C, and the aqueous phase was carefully transferred to a Phase Lock Gel (PLG) tube containing 600 µL of chloroform. After shaking to mix (without vortexing), the samples were centrifuged again for 5 minutes at 12,000 x g at 4°C. The aqueous phase was mixed with 0.5 mL of isopropanol and incubated at room temperature for 30 minutes at 4°C. RNA was pelleted by centrifugation at 16,000 x g for 10 minutes at 4°C, washed twice with 1 mL of cold 75% ethanol, and air-dried for 5-10 minutes. The RNA was resuspended in RNase-free water, and its concentration was measured using the RediPlate™ 96 RiboGreen™ RNA Quantitation Kit (Invitrogen) and diluted to 20 ng/µL for subsequent steps. RNA samples were stored at -80°C until further use. Library prep and sequencing was done by Novogene Co. Briefly, mRNA was purified from total RNA using poly-T oligo-attached magnetic beads. After fragmentation, the first strand cDNA was synthesized using random hexamer primers followed by the second strand cDNA synthesis. The library was ready after end repair, A-tailing, adapter ligation, size selection, amplification, and purification. The library was checked with Qubit and real-time PCR for quantification and bioanalyzer for size distribution detection. Quantified libraries were pooled and sequenced on NovaSeq 6000 with PE150 strategy, having ∼6GB of raw data per sample.

### Bioinformatics of RNA-seq data

Raw sequencing reads were quality filtered and adapter-trimmed using Trimmomatic with the following parameters: ILLUMINACLIP:TruSeq3-PE.fa:2:30:10, LEADING:3, TRAILING:3, and MINLEN:36^22^. These settings removed adapter sequences, trimmed low-quality bases (quality score <3) from both read ends, and discarded reads shorter than 36 bp after trimming. Filtered reads were aligned to the human reference genome (GRCh38; Ensembl release 113) using HISAT2^35^. Resulting SAM files were sorted by read name and converted to BAM format using samtools^36^. Gene-level counts from uniquely mapped reads were generated using featureCounts and combined to produce a count matrix across all samples^37^. Differential gene expression analysis was performed using DESeq2 with default parameters. Genes with an adjusted p-value (padj) < 0.05 and |log2FC| > 0.2 for Caco-2 cells or |log2FC| > 2 for macrophages were considered significantly up- or down-regulated. Hierarchical clustering was applied to both genes and samples using Euclidean distance with average linkage, and expression values were z-score normalized across genes before visualization. GO enrichment was performed using Enrichr^38,39^. Upstream regulator analysis was performed for differentially expressed genes using Ingenuity Pathway Analysis (IPA, QIAGEN Inc., https://digitalinsights.qiagen.com/IPA)

### Lipidomics and metabolomics sample preparation

For untargeted metabolomics, bEV pellets were resuspended in cold metabolite extraction buffer (acetonitrile:methanol:water, 2:2:1 [v/v/v]) prepared fresh with LC-MS grade solvents and Milli-Q 18 mΩ water, vortexed, and clarified through 0.22-µm Spin-X centrifuge filter tubes (VWR) by centrifugation. The resulting extract was dried and resuspended for LC-MS analysis. Metabolites were resolved by tandem reverse-phase and hydrophilic interaction liquid chromatography (HILIC) and detected on a high-resolution Orbitrap mass spectrometer^40^. For lipidomics, bEV pellets were extracted in 2:1 chloroform:methanol (v/v) with rotation for 24 hours at room temperature, then centrifuged at 4,000 RPM for 15 min at 4°C. The organic phase was decanted into a fresh Pyrex glass screw-top vial, evaporated using a GeneVac EZ-2 evaporator (Setting: 06-Low BP Mixture, 40°C), and rinsed with a minimal volume of 1:1 chloroform:methanol. The dried lipid residue was resuspended at 1 mg/mL in 1:1 chloroform:methanol and stored at −20°C. Prior to injection, 150 µL aliquots were evaporated and resuspended in 150 µL of solvent C (hexane:isopropanol, 70:30 [v/v], 0.02% formic acid, 0.01% ammonium hydroxide), centrifuged at 4,000 RPM for 15 min, and 100 µL of supernatant was loaded onto a Diol 3 µm Inertsil column (GL Sciences) for LC-MS analysis.

### Lipidomics and metabolomics analysis

Untargeted, pairwise lipidomics and metabolomics LC-MS data comparing cell pellets and bEVs were processed separately for positive- and negative-ionization modes using the XCMS Online platform^41^, following peak picking with MSConvert^42^. Post-acquisition intensity normalization was not applied, as equal amounts of total lipids or metabolites were injected for all samples prior to LCMS acquisition. Raw features were filtered using a minimum intensity threshold (≥ 5,000), statistical significance (p ≤ 0.05), and false discovery rate correction (q ≤ 0.05). Putative lipid and metabolite annotations were assigned using the LipidMaps and CEU Mass Mediator databases, respectively^43,44^. A lipid or metabolite species was classified as enriched in bEVs if it was detected in at least two bEV replicates and exhibited a higher mean intensity in bEVs than in cell pellets. Conversely, species were classified as depleted in bEV if they were detected in at least two cell pellet replicates and showed higher mean intensity in cell pellets than in bEVs. For features with zero signal in one condition, fold changes were calculated using the minimum intensity derived from the smallest non-zero observed value to avoid infinite ratios, according to the following formula:

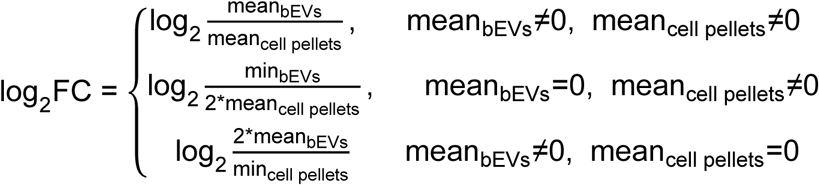

For downstream analysis, positive- and negative-ionization mode features sharing the same molecular formula were combined by summing replicate intensities across both modes prior to statistical comparison, ensuring that complementary ionization of the same species did not result in duplicate quantification. Mean bEV and cell pellet intensities were then recalculated from the summed replicates, and log₂ fold changes (log₂FC) were recomputed on the combined values using the same imputation strategy described above. Only metabolite annotations with at least one documented microbial association in the MiMeDB database were retained for further analysis, providing a biologically informed filter to exclude metabolites with no known bacterial relevance.

### Isolate-derived bEV proteomics sample preparation and analysis

OMV-containing samples (30 µL) were lysed in 4% SDS prepared in 50 mM ammonium bicarbonate, followed by freeze-thaw treatment, heating at 95 °C for 10 min, and probe sonication. After centrifugation (21,300 × g, 15 min), the supernatant was collected and protein concentration was measured with NanoDrop (ThermoFisher). Proteins were reduced and alkylated, followed by quenching of excess alkylating reagent with dithiothreitol. Samples were then processed using a modified single-pot, solid-phase-enhanced sample preparation (SP3) workflow with magnetic carboxylate beads (Cytiva)^45^. Proteins were captured by ethanol-induced precipitation, centrifuged, and washed three times with 80% ethanol, and digested overnight at 37 °C with trypsin/Lys-C at a 1:25 enzyme-to-protein ratio in 50 mM ammonium bicarbonate. Peptide-containing supernatants were recovered after magnetic bead separation, dried under vacuum, and stored at −20 °C until analysis. Peptides were resuspended in solvent A containing 6 x 5 LC-MS/MS Peptide Reference Mix (Promega) and injected into an Ultimate 3000 HPLC system coupled to an Orbitrap Exploris 240 mass spectrometer (MS). Peptides were separated on a reverse-phase C_18_ analytical column (Acclaim PepMap 100 2µm 75µm×150mm, ThermoFisher) at 30 °C using an 80 min gradient at a flow rate of 300 nL/min. MS was performed in positive-ion mode using data-dependent acquisition. Full-scan MS spectra were acquired over an *m/z* range of 400-1,600 at 60,000 resolution with a top 20 most intense precursor ion method for higher-energy collisional dissociation MS/MS selection. MS^2^ spectra were acquired at 30,000 resolution with dynamic exclusion of 100 s. Raw data were quality-checked in PReMiS software (Promega) using the reference peptide mix by assessing peak shape, retention-time stability, mass accuracy, and signal intensity. Database searching was performed in Proteome Discoverer 3.0 with Sequest HT against the relevant bacterial proteome and a contaminant database^46^. Searches were performed with full-tryptic specificity, up to two missed cleavages, precursor and fragment mass tolerances of 20 ppm and 0.05 Da, respectively, and a false discovery rate of 1%. Carbamidomethylation of cysteine was set as a fixed modification, whereas oxidation of methionine, deamidation, and common protein N-terminal modifications were included as variable modifications. Label-free quantification was based on peptide feature intensities, and relative protein abundance was estimated using the scaled total-ion current normalization approach, as shown in the equation below, where *I*’_j_ is the normalized intensity, *I*_j_ is the summed intensity of peptides assigned to protein *i*, *TIC*_all sample_ is the total summed intensity for each sample^47^.

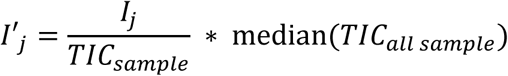

### Homolog and PPI prediction, and correlation analysis

Predicted protein structures were obtained from the AlphaFold database via UniProt when available. For proteins lacking precomputed models, structures were predicted using AlphaFold3 with default parameters and executed on the NSF Bridges-2 high-performance computing cluster^48,49^. Only predicted models with a predicted TM-score (pTM) ≥ 0.5 were retained for downstream analyses. To identify bacterial-human structural homologs, bacterial protein structures were aligned against human AlphaFold models using Foldseek^50^. Protein pairs covering atleast 80% of bacterial protein length with an alignment TM-score ≥ 0.5, e-value<10^-5^, and homolog probability (prob) = 1 were considered structurally homologous. For protein-protein interactions (PPIs), we compiled a database of human-commensal PPIs by integrating two complementary sources: structural homologs of experimentally validated human-bacterial PPIs, and disease-associated commensal proteins identified through metagenomic analysis. Candidate interactions were prioritized using AlphaFold3-based complex structure prediction. To assess the relationship between bEV proteome cargo and the transcriptomic response of Caco-2 cells, the UniRef50-based protein abundance matrix of bEVs and the log₂ fold-change transcriptome matrix of Caco-2 cells were log-transformed and z-score normalized. Pairwise Spearman correlations were computed, and protein-transcript pairs with |ρ| > 0.8 and p < 0.01 were considered significant. Within this significant set, transcription factors (TFs) were retained^51^, and their downstream targets were extracted from DoRothEA (regulons with confidence levels A, B, and C)^52–54^. Reactome pathway enrichment of these TF target sets was then performed in R using the ReactomePA^55^, with the full set of detected Caco-2 genes used as the background universe. Enriched terms with adjusted p < 0.05 and at least 5 overlapping genes were visualized.

## Results

### Diverse bacteria produce bEVs in the healthy human gut

To comprehensively characterize the diversity of bEVs present within the human gut, we collected stool samples from 11 healthy human subjects and isolated and purified stool-derived bEVs. We confirmed successful bEV isolation through negative staining transmission electron microscopy (TEM) (**Figure 1A**). We profiled the species and genes present in the stool samples through metagenomic sequencing (**Supplementary Figure 1A**). We then performed metaproteomics to identify the taxonomic sources of bEVs. Our results show that the predominant protein intensities (at the label-free quantification (LFQ) and peptide-level) were derived from bacterial sources (**Figure 1B**). The most prevalent bEVs were produced by bacteria in the phyla Bacteroidetes (recently renamed Bacteroidota) and Firmicutes (recently renamed Bacillota), prominently featuring well-known gut commensals such as *Faecalibacterium prausnitzii* and *Phocaeicola vulgatus* (**Figure 1C, Supplementary Figure 1B**). There was greater inter-personal variation in certain bEVs, such as those produced by bacteria belonging to genera such as *Bacteroides*, *Segatella*, and *Bifidobacterium*. The abundances of bEVs correlated to some extent with their cognate bacterial abundances (**Supplementary Figure 1C**). While the diversity of vesiculating bacteria remains debated^56,57^, our findings suggest that vesiculation among human gut bacteria is far more widespread than previously appreciated.

**Figure 1.**
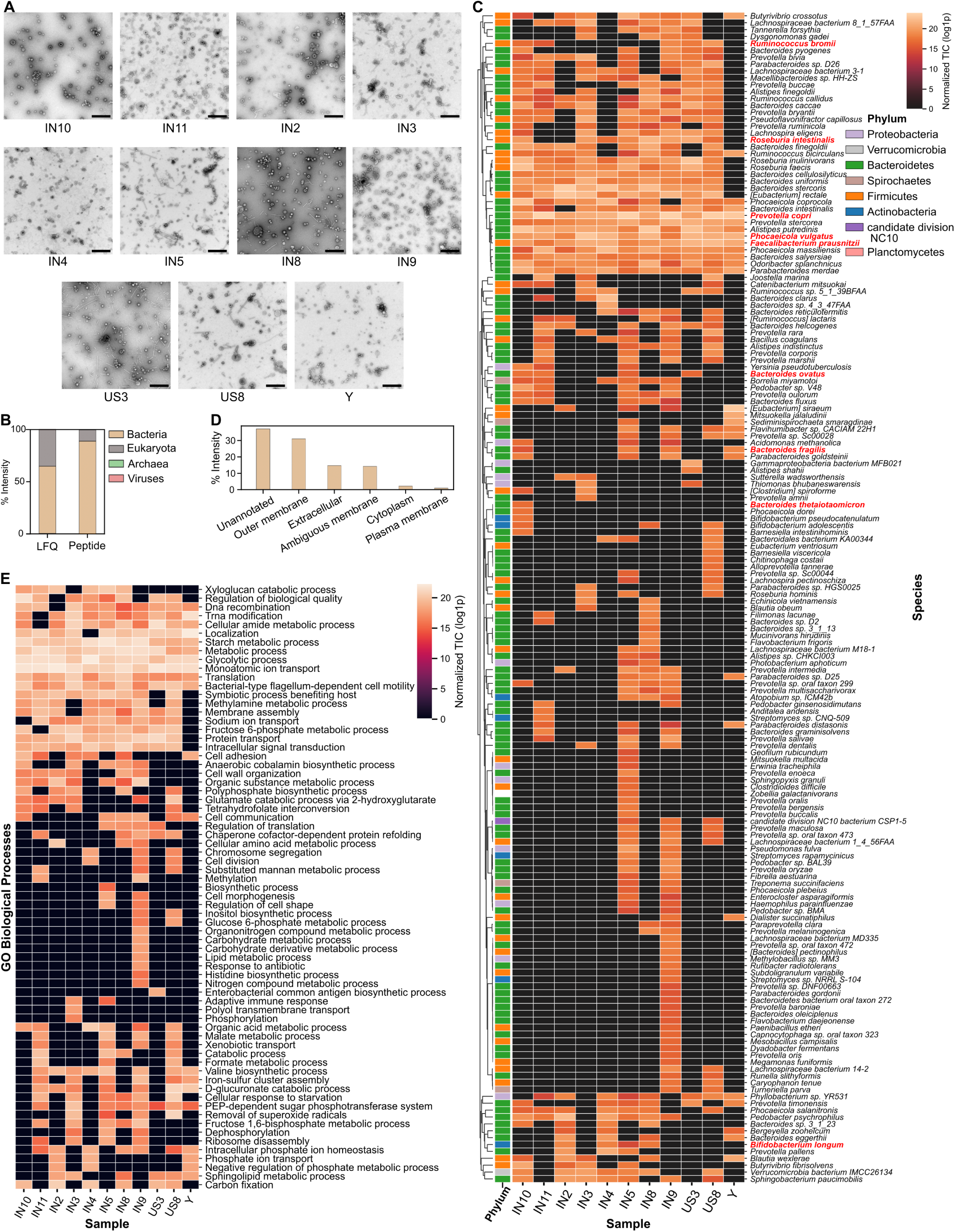
Taxonomic and functional landscape of bEVs isolated from human stool. (A) Representative TEM images of purified bEVs isolated from human stool samples (scale bar, 500 nm). (B) Relative contribution of protein intensity from different kingdoms at the LFQ and peptide levels from combined proteomics across all human stool samples. (C) Taxonomic distribution of bEV-associated proteins at the species level. Colors represent normalized TIC (total ion count) values. Side color bar represents phyla. Species names appearing in red are those from which bEVs were isolated in later experiments. (D-E) Collapsed gene ontology (GO) terms of proteins identified in stool-derived vesicles, shown for Biological Process (D) and Cellular Component (E) categories. Color represent normalized TIC values.

The majority of annotated proteins are predicted to localize predominantly to the bacterial outer membrane, although we also find a large portion of extracellular and cytoplasmic proteins (**Figure 1D**). Notably, proteins annotated as localizing to the plasma membrane were relatively scarce. We attribute this to current limitations in protein annotation methods, which are particularly limited when annotating inner membrane or periplasmic proteins. To characterize the functional capabilities and biological significance of these bEVs, we examined the Gene Ontology (GO) annotations of the identified vesicular proteins. Notably, bEV proteins were annotated into various biological processes, including proteolysis, general metabolic activities, monoatomic ion transport, and components of the tricarboxylic acid (TCA) cycle (**Figure 1E**). There were also proteins annotated with specific host-interaction functions. These included microbial lipoproteins and flagellins from diverse taxa that are known to interact with host immune receptors such as TLR2/TL4/TLR5 (**Supplementary Table 1**). We also detected proteins with previously described roles in gut homeostasis or host modulation, including the microbial anti-inflammatory molecule (MAM) from *F. prausnitzii*, which has been shown to inhibit IL-1β-induced IL-8 secretion in intestinal epithelial Caco-2 cells and promote gut barrier integrity^58^. In contrast, several proteases, primarily from Bacteroidota, were also present, including M60-like proteases which are known to degrade mucin and potentially remodel extracellular matrices^59^. Additional host-interacting proteins included SusD/RagB-family proteins from *Bacteroides fragilis*, which can bind to the host NOD1 receptor and enhance chemoresistance^60^, and OmpA-like proteins from *Parabacteroides goldsteinii*, previously reported to induce IL-22 signaling^61^. Together, these findings suggest that stool-derived bEVs carry a diverse set of proteins capable of directly interacting with host cellular pathways.

### Gut bEVs exhibit diverse phenotypes and host interactions

Based on the combined metaproteomic and metagenomic data, we selected nine representative bacterial species from which to isolate bEVs for in-depth analysis (**Figure 1C**). The selection was based on bacterial abundance across stool samples, choosing representatives of both high and low abundance, as well as taxonomic diversity. The selected isolates included *Bacteroides fragilis*, *Bacteroides ovatus*, *Bacteroides thetaiotaomicron*, *Phocaeicola vulgatus*, *Prevotella copri* (recently renamed *Segatella copri*), *Bifidobacterium longum*, *Faecalibacterium prausnitzii* (recently renamed *Faecalibacterium duncaniae*), *Roseburia intestinalis*, and *Ruminococcus bromii*. Due to difficulties culturing *R. bromii*, we substituted it with *Ruminococcus gnavus* (recently renamed *Mediterranean gnavus*), a closely related species. Additionally, we incorporated three known pathobionts, *Fusobacterium nucleatum*, *Enterococcus faecalis*, and *Streptococcus mutans* for comparison.

Given the limited prior studies reporting bEV isolation from *P. copri*, *R. intestinalis*, and *R. gnavus*, we initially performed scanning electron microscopy (SEM) on overnight bacterial cultures to confirm that these species do, in fact, produce bEVs under laboratory culture conditions. SEM revealed prominent surface blebbing across these strains, a hallmark of bacterial vesiculation^62^ (**Figure 2A**). Subsequently, we isolated and purified bEVs from these three species in addition to nine mentioned above and validated the presence of intact bEVs via TEM (**Figure 2B**). Consistent with expectations, Gram-positive bacteria produced single-membrane bEVs (IMVs/CMVs), while Gram-negative bacteria exhibited mostly single-membraned vesicles (OMVs) with some double-membraned vesicles (OIMVs), except for *B. fragilis* and *P. copri*, which predominantly generated OIMVs. bEVs varied by mean bEV diameter and size heterogeneity across different bacterial species (**Figure 2C**). However, within the *Bacteroides* genus, no significant differences in bEV size were detected.

**Figure 2.**
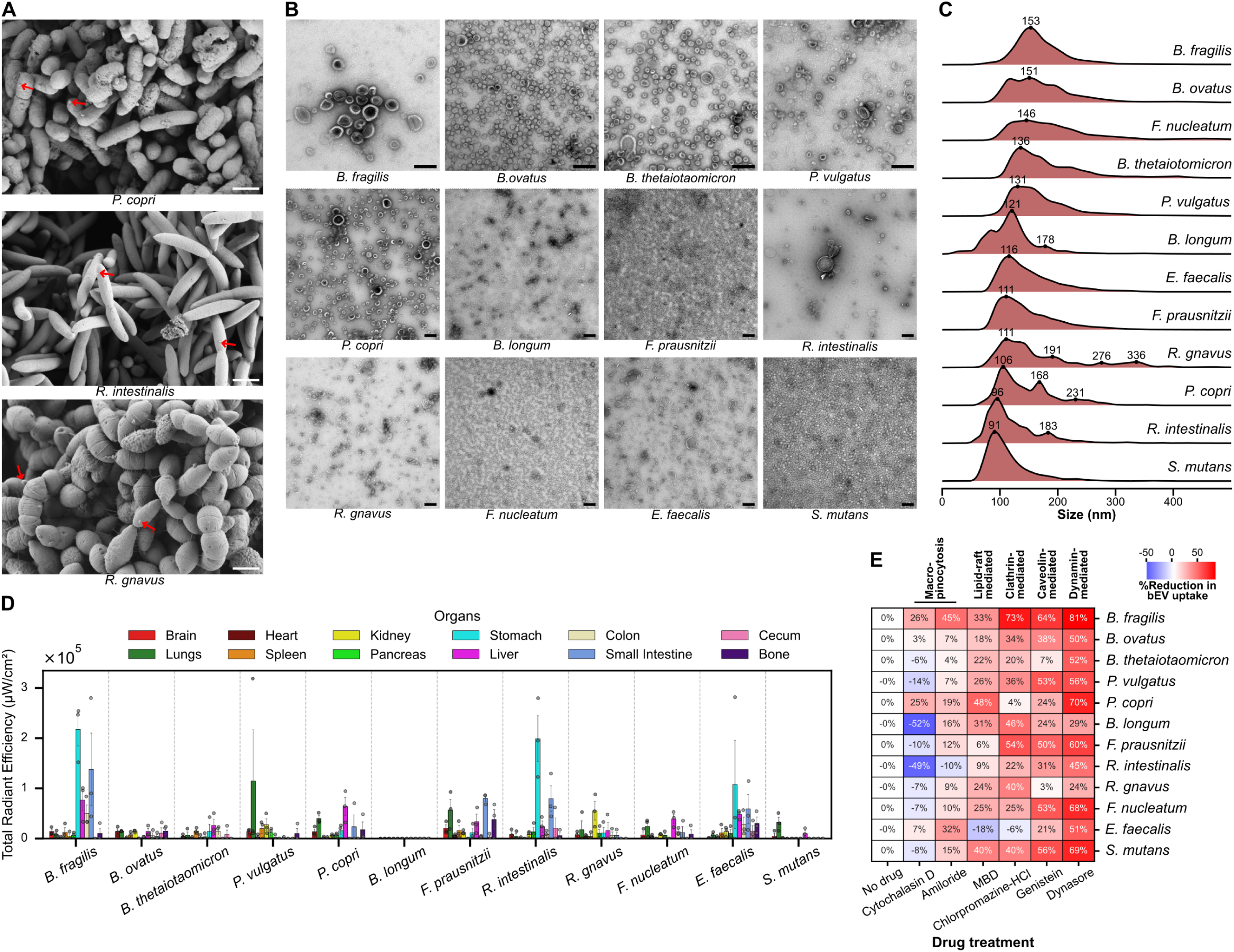
Characterization of isolate-derived bEVs and their endocytic uptake by epithelial cells. (A) SEM images of overnight bacterial cultures. Red arrows highlight membrane protrusions consistent with vesicle formation. Scale bar: 1 pm. (B) Representative TEM images of negatively stained, purified bEVs derived from 12 human gut bacterial species. Scale bar: 200 nm. (C) Nanoparticle tracking analysis (NTA)-based size distribution profiles of bEVs shown in (B). Numbers represent local peak values. (D) Barplot showing intensity and standard deviation of Cy7-labelled bEVs in organs from mice gavaged with 25pg (total protein) of bEVs from each species. Intensity values were normalized with organ weight and subtracted from the background (PBS-treated mice) to remove autofluorescence. (n=3 mice per condition) (E) Quantification of bEV uptake by flow cytometry following inhibition of endocytic pathways. Positive values indicate the percent reduction in uptake, whereas negative values indicate an increase in uptake relative to untreated controls. MBD: methyl beta-cyclodextrin.

Previous work shows that certain bEVs can translocate to extraintestinal body sites^1,2,63^. Therefore, we next sought to determine whether isolate-derived bEVs exhibited organ-specific distribution *in vivo*. We examined their biodistribution across 12 mouse organs (brain, lungs, heart, spleen, kidney, pancreas, stomach, liver, colon, small intestine, cecum, and bone) after oral gavage. Overall, bEVs preferentially localized to distinct tissues (**Figure 2D**). Consistent with their intestinal origin, many, albeit not all, isolate-derived bEVs accumulated within the gastrointestinal (GI) tract, despite extensive wash steps, similar to previous reports of *Akkermansia muciniphila* bEVs^63^. It should be noted, however, that we cannot distinguish whether the observed fluorescence signal reflects cellular internalization, or bEVs retained in the extracellular space or intercellular compartments of the tissue. *B. fragilis*-derived bEVs localized predominantly to the GI tract and liver, suggesting potential involvement in gut-liver axis communication. Others preferentially accumulated in extraintestinal tissues: *P. vulgatus* and *F. prausnitzii* in lung, *R. gnavus* in kidney, and *E. faecalis* in bone. The latter observation is noteworthy given the connection between *E. faecalis* and colorectal cancers (CRC) and may point towards a potential role in bone-related metastasis or CRC-associated bone loss^64,65^. We did not detect accumulation of *B. longum* bEVs in any organ examined, which may reflect a lack of tissue targeting, but may also be caused by technical factors, such as lower labeling efficiency, bEV stability, rapid clearance, or limitations in detection sensitivity. Additionally, we observed considerable mouse-to-mouse variability in bEV tissue localization, underscoring the need for further investigation. Taken together, these findings represent a critical first step in characterizing the organ-level distribution of gut bacteria-derived bEVs *in vivo* and lay the groundwork for future studies that will more precisely define their cellular fate, tissue targeting mechanisms, and role in systemic manifestations of gut-associated diseases.

We next asked whether these bEVs entered host cells and, if so, by what mechanisms. We determined that bEVs are non-cytotoxic to human intestinal epithelial cells (Caco-2 cells) (**Supplementary Figure 2A**). We examined the endocytic pathways by which these bEVs were taken up by Caco-2 cells by employing inhibitors specific to six different endocytosis mechanisms: cytochalasin D (blocks F-actin polymerization), amiloride (inhibits macropinocytosis), methyl-beta-cyclodextrin (disrupts lipid rafts), chlorpromazine-HCl (inhibits clathrin-mediated endocytosis), genistein (blocks caveolin/lipid raft-mediated pathways), and dynasore (inhibits dynamin-dependent endocytosis). Caco-2 cells pretreated with endocytosis inhibitors were exposed to DiD-labelled bEVs for 4 hours. We found that dynamin-mediated endocytosis was the most commonly utilized pathway across bEVs, although multiple uptake routes were involved for each bEV (**Figure 2E**). For example, bEVs from *Bacteroides* species primarily entered via dynamin-dependent endocytosis, except for *B. fragilis*, whose bEVs showed nearly equal reliance on dynamin-, clathrin-, and caveolin/lipid raft-mediated pathways for uptake. Across all organisms, methyl-β-cyclodextrin (MBD) produced modest decreases (6 % to 48%) in bEV uptake, suggesting that lipid rafts facilitate, but do not dictate, initial bEV- host cell membrane interaction or downstream signaling prior to endocytosis. Interestingly, cytochalasin-D, which inhibits actin polymerization, increased uptake (∼50% increase for *B. longum* and *R. intestinalis* (**Supplementary Figure 2B**)), suggesting that bEV entry is largely independent of actin-driven engulfment and may be favored by reduced cortical tension. Multiple studies have shown a similar increase in cytochalasin D-mediated increase in uptake, but the exact mechanism is still unknown^66–68^. These findings show that bEVs employ multiple routes of uptake, each of which can result in different cargo trafficking^69^.

### bEVs evoke differences in transcriptional responses from macrophages and intestinal epithelial cells

Macrophages are highly phagocytic innate immune signaling cells that surveil the intestinal epithelium and are therefore expected to engage bEV cargo through fundamentally different sensing and signaling pathways than epithelial cells^70^. To assess whether bEVs induce host responses and whether these were generalizable across cellular contexts, we next profiled the transcriptomes of intestinal epithelial cells and macrophages following bEV treatment (**Figure 3A**). By examining differentially expressed genes (DEGs) in the bEV-treated groups, we observed markedly divergent patterns between Caco-2 epithelial cells and macrophages, suggesting cell-type-specific transcriptional organization of bEV responses (**Figure 3B**, **Supplementary Figure 3A**). Transcriptional responses to bEVs clustered into four groups in epithelial cells and three in macrophages, each associated with a set of distinct species. However, we find a set of genes upregulated across all clusters in epithelial cells, and a second gene set across macrophages, indicating a shared core host response to bEVs (**Supplementary Figure 3B**).

**Figure 3.**
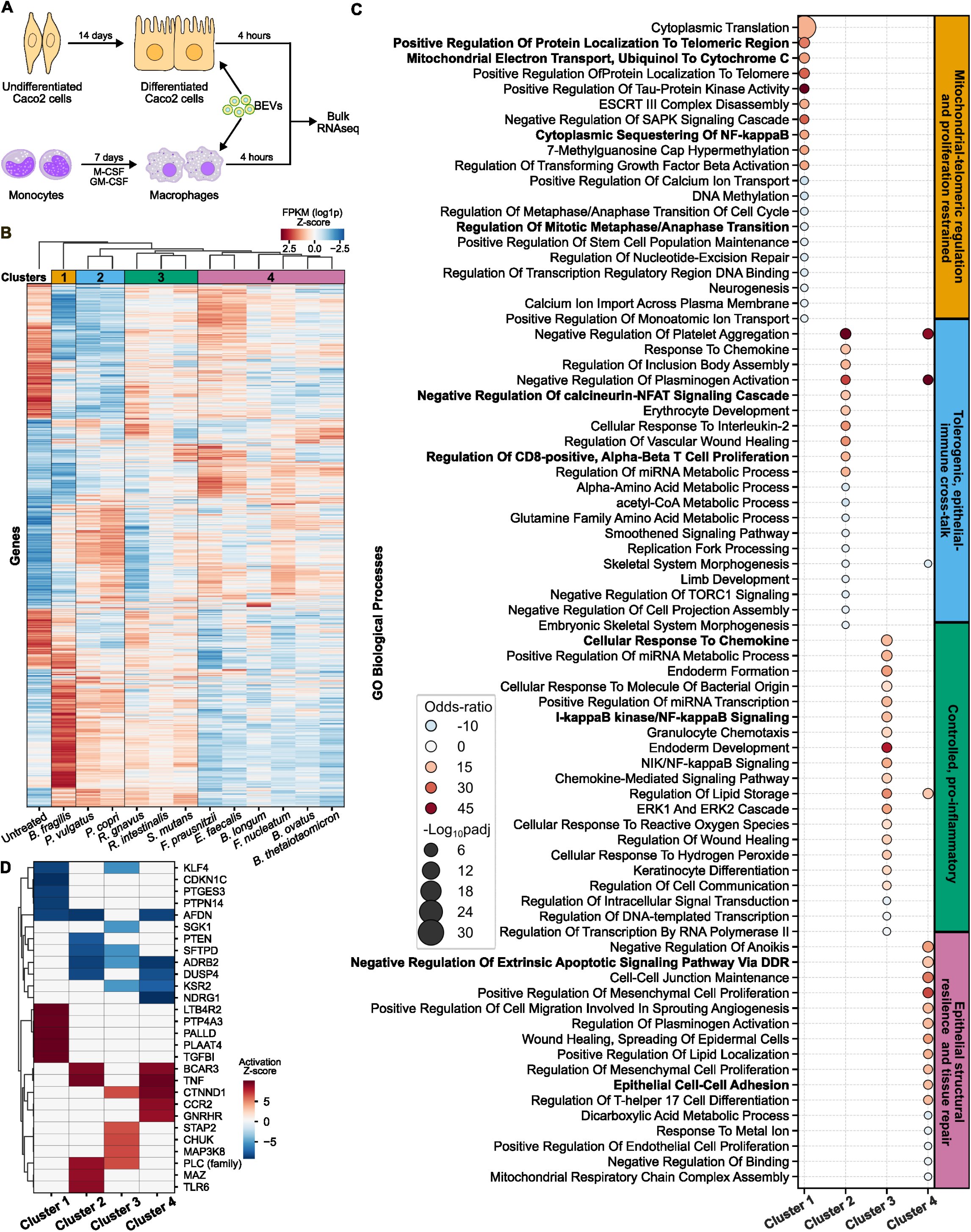
Transcriptomic profiling of host cells exposed to bEVs. (A) Schematic overview of the experimental design used to investigate transcriptomic responses to bEV. (B) Hierarchical clustering of samples based on FPKM-normalized (log) gene expression following treatment with bEVs in Caco-2 cells. Color represents z-scores by row. (C) Top 12 significantly enriched upregulated and downregulated Gene Ontology Biological Process (GO BP) pathways per cluster defined in (B). Color represents the odds-ratio, and size is the log(adjusted p-value). (D) Upstream regulator analysis performed using Ingenuity Pathway Analysis (IPA) to identify key transcriptional drivers of the observed per-cluster responses. Color represents the z-score of activation. A positive z-score signifies activation, and a negative value signifies repression.

Focusing on epithelial cells, we next characterized the molecular terms (defined by GO-BP^39^) defining each cluster. Cluster 1, containing bEVs solely from *B. fragilis*, was characterized by transcriptional signatures related to mitochondrial-telomeric regulation with anti-inflammatory and proliferative restraint^71^. Additionally, we observe increased translational activity, suggesting a shift in resources from cell division to host defense. Cluster 2, defined by bEVs from *P. copri* and *P. vulgatus*, reflected a tolerogenic (TGF-β1-mediated) and increased epithelial-immune crosstalk, indicated by cytokine and chemokine responses. Cluster 3, driven by bEVs from *R. gnavus, R. intestinalis, and S. mutans*, all Gram-positive organisms, showed fewer total DEGs (202 upregulated and 76 downregulated genes) compared to more than 1,000 DEGs in other clusters, hinting towards suppressed/controlled response. Of the active genes, they were enriched in pro-inflammatory pathways. In contrast, Cluster 4, represented by *E. faecalis, F. prausnitzii, B. longum, F. nucleatum, B. ovatus*, and *B. thetaiotaomicron* bEVs, displayed a transcriptional profile centered on epithelial survival and active tissue repair, indicated by the upregulation of anoikis resistance, cell-cell junction maintenance, and epithelial adhesion, consistent with commensal-rich groups’ known ability to promote tissue repair and cell survival^72,73^.

We next sought to identify regulatory targets of bEV-induced epithelial responses. We used Ingenuity Pathway Analysis^74^ to identify upstream regulators of the transcriptional responses observed in Caco-2 epithelial cells driving cluster-specific responses (**Figure 3D**). Predicted master regulators recapitulated many of the biological themes identified from GO BP analysis of DEGs. Of note, Cluster 1 is regulated by PTGES3. Although this protein normally functions as a prostaglandin synthase, it also interacts with HSP90 and hTERT to promote telomerase activity^75^, characterized by DEGs in this cluster. Given that Caco-2 cells retain telomerase activity as a property of their cancer cell origin, this signal may reflect modulation of an inherent cancer cell survival program rather than a response typical of normal intestinal epithelium. In Cluster 2, TNF activation drives immune-epithelial communication (via IL-2, CD8+, chemokine signaling). In Cluster 3, the regulatory picture that emerged was one of disinhibited NF-κB signaling. We observe suppression of KLF4, which regulates goblet cell differentiation and mucin production in the colon^76^; and activation of CHUK and MAP3K8, which directly engaged upstream NF-κB kinase machinery, together producing the controlled pro-inflammatory state we observe. In Cluster 4, activation of CTNND1, which stabilizes E-cadherin-mediated junctions and negatively regulates Wnt signaling, a pathway central to intestinal crypt homeostasis and epithelial renewal^77^, directly supported the junctional integrity and epithelial survival programs enriched in this cluster.

In macrophages, although three clusters were identified, transcriptional responses showed greater overlap across bEV groups than epithelial cells (**Supplementary Figure 3C**). Shared terms included regulation of viral entry pathways, responses to viral and bacterial components, and activation-associated programs such as cytokine and chemokine signaling, with differences largely reflecting variation in magnitude rather than distinct biological programs. Cluster-specific features were comparatively subtle: Cluster 1 was enriched for metal ion regulation and protein ubiquitination pathways, Cluster 2 for IL-15-associated phagocytic signaling, and Cluster 3 for anti-inflammatory IL-6 responses^78^. In contrast to the more diverse and functionally segregated programs observed in Caco-2 epithelial cells, macrophage responses were more convergent, capturing a shared early innate immune activation state.

### bEVs carry diverse molecular cargo that interacts with cellular transcriptional modules

To assess whether the distinct transcriptional responses elicited by bEVs could be attributed to differences in their molecular cargo, we first profiled each isolate’s bEV cargo, using proteomics, lipidomics, and metabolomics. Lipidomic profiling revealed that bEVs and cell pellets were predominantly composed of glycerophospholipids, glycerolipids, and sphingolipids across all strains, but the compositional differences at this level were modest in magnitude (**Supplementary Figure 4A**). A notable exception was glycerophosphoserine (PS), which was significantly enriched in bEVs across all strains except *B. ovatus* and *F. prausnitzii*; PS enrichment on the bEV surface may facilitate host cell entry by mimicking apoptotic bodies and engaging TIM/TAM receptor-mediated endocytosis^79^. Fatty acyl lipids were enriched in bEVs from most Gram-negative bacteria, but depleted in Gram-positive-derived bEVs, where they constituted over 50% of the detected lipid classes in cell pellets. In *P. copri*, sterol lipids accounted for approximately 50% of the total lipid composition, dominated by an annotation putatively identified as 25-hydroxyvitamin D2 25-(beta-glucuronide), which has not been previously reported in bacteria.

Metabolomic profiling further revealed bEV-specific packaging patterns (**Supplementary Figure 4B)**. 5-Hydroxyisourate, an intermediate in bacterial purine catabolism, was enriched in bEVs from *B. fragilis*, *P. vulgatus*, and *B. longum* (log₂FC 2.6-3.5), consistent with its known presence in Bacteroidetes and Firmicutes. The most broadly depleted metabolites across bEVs compared to cell pellets were *N*-acyl amino acids, including linoleamide, *N*-myristoyl lysine, *N*-oleoyl arginine, and *N*-undecanoyl glycine, suggesting that there is selective exclusion of these membrane-associated aminolipids during bEV biogenesis or that these aminolipids do not facilitate vesiculation.

As expected, bEV proteomes clustered according to phylogeny (**Supplementary Figure 5A**). Gene ontology terms showed that enzymes involved in carbohydrate, lipid, and amino acid metabolism were consistently represented across nearly all bEV populations, reflecting the packaging of central metabolic machinery into bEVs (**Supplementary Figure 5B**). Beyond this shared metabolic core, however, species-specific cargo profiles pointed to distinct functional roles in host-microbiome interactions. Bacteroidota-derived bEVs were abundant in fimbrillins, which facilitate bacterial adhesion to mucosal surfaces^80^ where it may aid in epithelial cell uptake (**Figure 4A, Supplementary Table 2**). Additionally, iron acquisition machinery was also prominently represented in the *Bacteroides* bEVs, which has been previously reported to be exploited by bystander organisms^81^. Elongation factor Tu (EF-Tu), while canonically a translation-associated GTPase, was detected across all 12 bEV populations and ranked among the most abundant protein cargoes in the Gram-positive species *R. intestinalis*, *R. gnavus*, *E. faecalis*, and *B. longum.* EF-Tu is also well established as a moonlighting protein, relocating to the bacterial cell surface where it interacts and adheres to host cells^82^. EF-Tu is also a recognized activator of innate immune signaling in epithelial cells and immune cells, likely through TLRs, driving the production of pro-inflammatory cytokines such as IL-8, TNF-α, and IL-6, and promoting dendritic cell maturation^82^. It can also interact with the intracellular domain of CD21^83^. Its conserved presence in bEVs across phylogenetically diverse commensals suggests that bEV-delivered EF-Tu may help establish tonic immune activation levels, contributing to the balanced surveillance that maintains homeostasis at the gut mucosal surface. In *F. nucleatum*-derived bEVs, we find FadA and Fap2, which, when present in bEVs, have been shown to facilitate bacterial colonization in colorectal cancer tissue and exacerbate inflammation at distal sites, including synovial joints^2,7^. *S. mutans* bEVs were overwhelmingly dominated by glucosyltransferase-I (GTF-I), its principal virulence factor. GTF-I-harboring bEVs have been shown to inhibit biofilm formation of competing oral Streptococci^84^. Beyond their capacity to modulate host transcription, bEVs may also engage host cells through direct protein-protein interactions. Indeed, we found that bEVs across multiple species carried proteins predicted to interact with human proteins or mimics of human proteins, although many of these have yet to be experimentally validated (**Figure 4A**). These findings together highlight an underexplored reservoir of potential molecular mediators in host-microbiome crosstalk.

**Figure 4.**
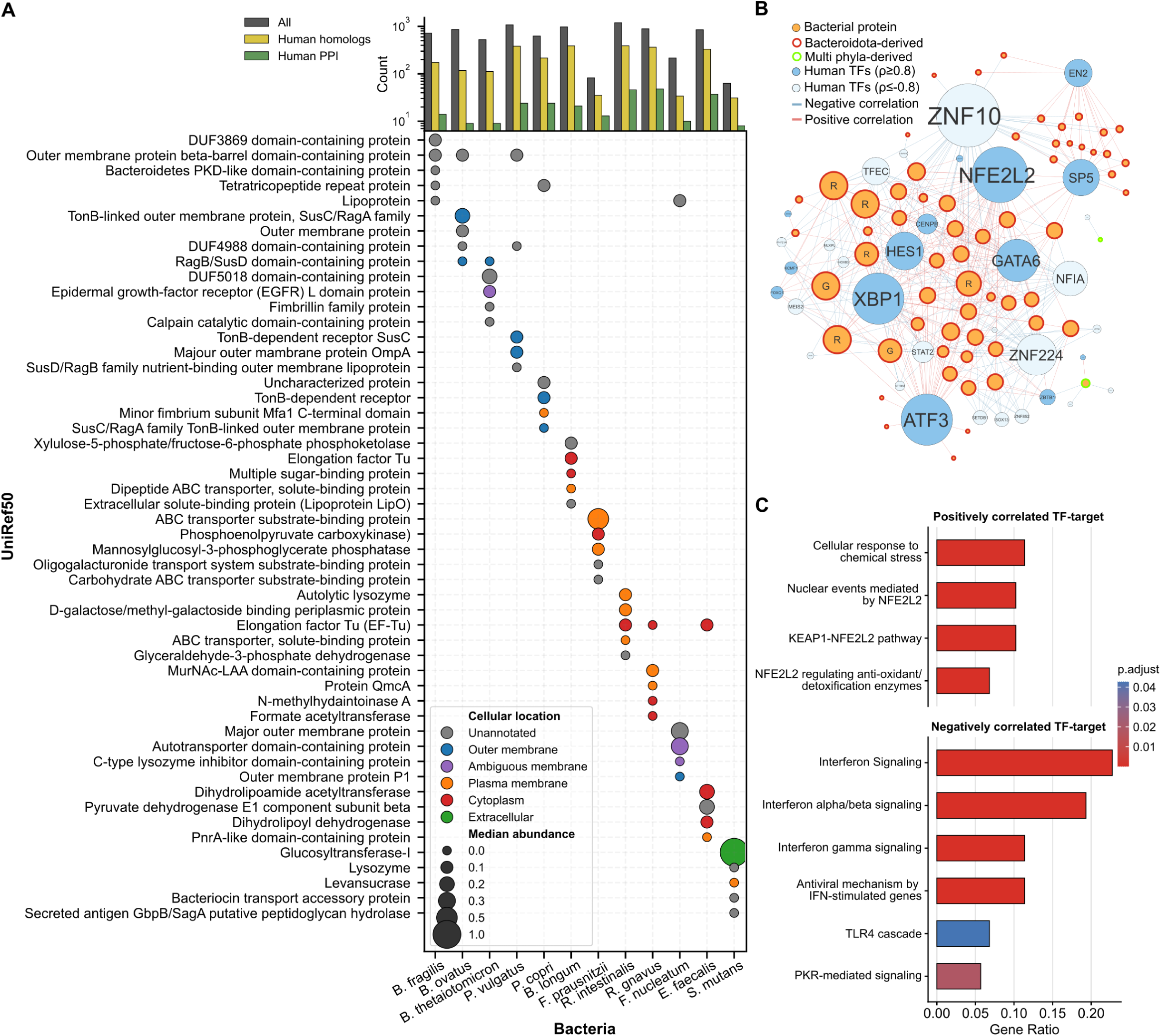
Integration of the bEV proteome with epithelial transcriptomic responses. (A) The five most enriched UniRef50 functional categories across each bEV proteome from 12 bacterial species. Colors represent the cellular location; size represents median abundances. The upper panel represents the number of bacterial proteins detected (gray), proteins with shared structural homology with human proteins (yellow), and proteins with predicted interactions with human proteins (green). (B) Network visualization of significant proteome-transcriptome correlations between bEV cargo (orange) and Caco-2 transcription factors (TFs) (blue). Node and font size reflect the number of connections (degree). R: bacterial ribosomal proteins; G: bacterial glycolytic proteins. (C) Reactome pathway enrichment analysis of targets regulated by TFs correlated with the highest-degree bacterial proteins. Color represents the adjusted p-values.

To investigate which bEV protein cargo might be driving the transcriptional response in Caco-2 cells, we performed a correlation analysis. We focused our analysis on human transcription factors (TFs) as they would capture downstream signaling more comprehensively than individual effector genes. We identified 288 strong correlations (|ρ| ≥ 0.8, p < 0.01) linking 62 bacterial proteins to 34 human TFs (**Supplementary Table 3**, **Figure 4B)**. All TFs consistently responded to bacterial proteins, correlating positively or negatively. Positively corelated TFs included the stress-responsive TFs ATF3, NFE2L2 (Nrf2), and XBP1, alongside the intestinal differentiation regulators GATA6, HES1, SP5, and EN2 (**Supplementary Figure 4C**). ZNF10 was the most negatively connected TF in the network, with 29 bacterial protein partners, along with ZNF224, NFIA, and TFEC. However, some of these TFs are co-expressed and/or act within the same transcriptional module, explaining their similar responses to the same sets of bacterial proteins (**Supplementary Figure 5D**). Bacteroidota-derived ribosomal proteins and glycolytic enzymes were highly represented. These Bacteroides-derived bacterial proteins collectively were involved in NFE2L2-driven cytoprotective and antioxidant pathways and negatively correlated with interferon signaling (**Figure 4C**), suggesting that Bacteroidota bEV cargo activates epithelial stress defense while dampening interferon-mediated immune signaling.

## Discussion

bEVs represent an underappreciated mechanism by which microbes can communicate with host tissues and influence microbiome-related disorders. In this study, we examined vesiculation in human stool samples, identifying a diverse taxonomy of bEV producers. We also evaluated the capacity of bEVs to reach tissues beyond the gut, and applied integrated multi-omics approaches to characterize their molecular cargo and functional impact on host cells. Together, our findings reveal both conserved and cell-type-specific transcriptional programs elicited by bEV exposure, highlighting a structured framework of bEV-mediated microbiome-host communication.

Whether vesiculation is a universal feature of bacteria has long been debated. Although numerous *in vitro* studies have shown that environmental conditions such as temperature^85^, stress^86^, and antibiotic exposure^87^ can substantially influence bEV production, our findings suggest that vesiculation may be far more widespread among gut bacteria *in vivo* than previously appreciated. By directly examining human stool-derived samples, we demonstrate that a broad range of commensal species release bEVs under physiological conditions, indicating that bEV production is not a rare phenomenon but rather a common trait within the healthy gut ecosystem. Importantly, our data also indicate that bacterial abundance alone may not predict bEV-mediated impacts *in vivo*. Differences in vesiculation rates across species imply that low-abundance bacteria could exert disproportionate influence through high bEV production, whereas highly abundant taxa may contribute fewer bEVs.

Beyond their prevalence, our multi-omics characterization reveals that bEVs carry diverse molecular cargo, including proteins with predicted host-interaction potential. Importantly, bEVs are not inert byproducts; instead, our data support that bEV cargo induces measurable transcriptional changes in recipient host cells. bEVs carry known immunomodulatory cargo such as EF-Tu^82^, OmpA^61^, and MAM^58^, that contribute to tolerogenic or pro-inflammatory responses. bEV cargo can engage with host tissues either on the surface or can escape the endosome once internalized. Gram-negative derived bEVs are coated with LPS and FomA, a surface protein on *F. nucleatum-*derived bEVs, are agonist of surface receptors TLR4 and TLR2^88^, respectively. On the other hand, OmpA from *A. baumannii* bEVs leaks into the host cytoplasm to cause mitochondrial fragmentation-related cytotoxicity^11^ and internalized LPS from EHEC activates cytoplasmic caspase-mediated pyroptosis^10^.

Although not the central focus of this work, bEVs also play important community-shaping roles. The proteomic landscape of bEVs, abundant in SusC/SusD polysaccharide utilization components, glycoside hydrolases, and nutrient-binding lipoproteins, that we observe is consistent with prior work showing that *Bacteroides*-derived bEVs can degrade dietary and host glycans, with breakdown products serving as “public goods” that support the growth of other community members^89^. Furthermore, the detection of fimbrillins in Bacteroidota bEVs, virulence-associated adhesins such as FadA in *F. nucleatum* bEVs, and glucosyltransferase-I in *S. mutans* bEVs participate in colonization, niche competition, and biofilm formation.

The observations that epithelial cells actively internalize bEVs, that bEVs contain potential molecular mimicry or host-interacting proteins, and that bEV cargo strongly correlates with host transcriptional programs raise fundamental evolutionary questions. It may be that the host has evolved mechanisms to sense and internalize bEVs as part of mucosal surveillance, supported by increasing observations that germ-free animals exhibit impaired mucosal immune development that can be partially rescued by bEV administration alone^12,63,90^. Alternatively, bEVs possess molecular features, such as glycerophosphoserines, that have been selected to facilitate host uptake, suggesting microbial adaptation for interkingdom communication. Or it may be a combination of both. Our transcriptomic analyses further reveal that bEV-induced responses are both bEV-origin specific and cell-type dependent, highlighting functional specialization in microbiome-host communication. *B. fragilis* bEVs have been shown to promote anti-inflammatory effects on immune cells^12^, a finding that aligns with the suppression of inflammatory signaling and proliferative restraint we observed in *B. fragilis* bEV-exposed intestinal epithelial cells. We broaden the role of *B. fragilis* bEVs to include effects on mitochondrial gene expression and telomeric maintenance. Similarly, our findings directly corroborate previous observations that prevalent commensals regulate tight junction gene expression in epithelial cells via bEVs^72,91^. Beyond their epithelial effects, bEVs present in the gut lumen can cross the epithelial barrier to disseminate systematically, suggesting that the transcriptional states we describe here may represent only the first layer of a broader, multi-cellular host response to commensal bEV exposure.

This is the first investigation into the sources of stool-derived bEVs via metagenomics and metaproteomics. Our study focused on bEV production in healthy individuals to establish a baseline of the healthy gut, although our focus will expand in the future to include bEV production and distribution in inflammatory disease conditions, where the abundances of organisms and other signaling cues are different^92^. To gain better resolution of bEV cargo, we isolated bEVs from 12 gut bacteria cultured under laboratory conditions, employing the most stringent isolation protocols, however laborious, to ensure the purity of our bEV preparations. This allowed us to comprehensively characterize bEV-associated proteins, lipids, and metabolites. bEVs also carry nucleic acids, which may contribute to host modulation and warrant future investigation^93^. Despite studying isolate-derived bEVs, bEV production and cargo composition in the native gut environment may differ due to host-derived signals, microbial interactions, and spatial niche effects. Additional species- and strain-level variation may further diversify bEV cargo composition and function. Finally, studying bEVs *in vivo* presents technical challenges, including difficulties in isolating endogenous bEVs from complex tissues and accurately quantifying physiological exposure levels. Continued methodological advances will be important to refine our understanding of bEV abundance, distribution, and functional impact *in situ*.

## Data and code availability

All the data generated in the manuscript will be available after publication.

## Acknowledgements

This study is supported by the Packard Foundation and NIH grant (1R01LM014714-01A1). We thank the Proteomics Resource Center at Rockefeller University (RRID: SCR_017797) for stool bEV proteomic analyses, performed using instrumentation funded by the Sohn Conferences Foundation and the Leona M. and Harry B. Helmsley Charitable Trust. We also acknowledge the Cornell Flow Cytometry Core Facility (RRID: SCR_021740), Cornell Imaging Facility (NIH S10OD025049), and the Cornell Center for Materials Research (CCMR) for providing access to imaging and flow facilities, and the staff at the Microscopy and Image Analysis Core at Weill Cornell Medicine for assistance with electron microscopy. We are grateful to the Department of Molecular Medicine (College of Veterinary Medicine), the Baker Institute for Animal Health, and the Weill Institute for Cell and Molecular Biology at Cornell University for access to ultracentrifugation facilities, and to the Cerione laboratory at the College of Veterinary Medicine for access to NanoSight NTA instrumentation.

---

**Supplementary Figure 1.**
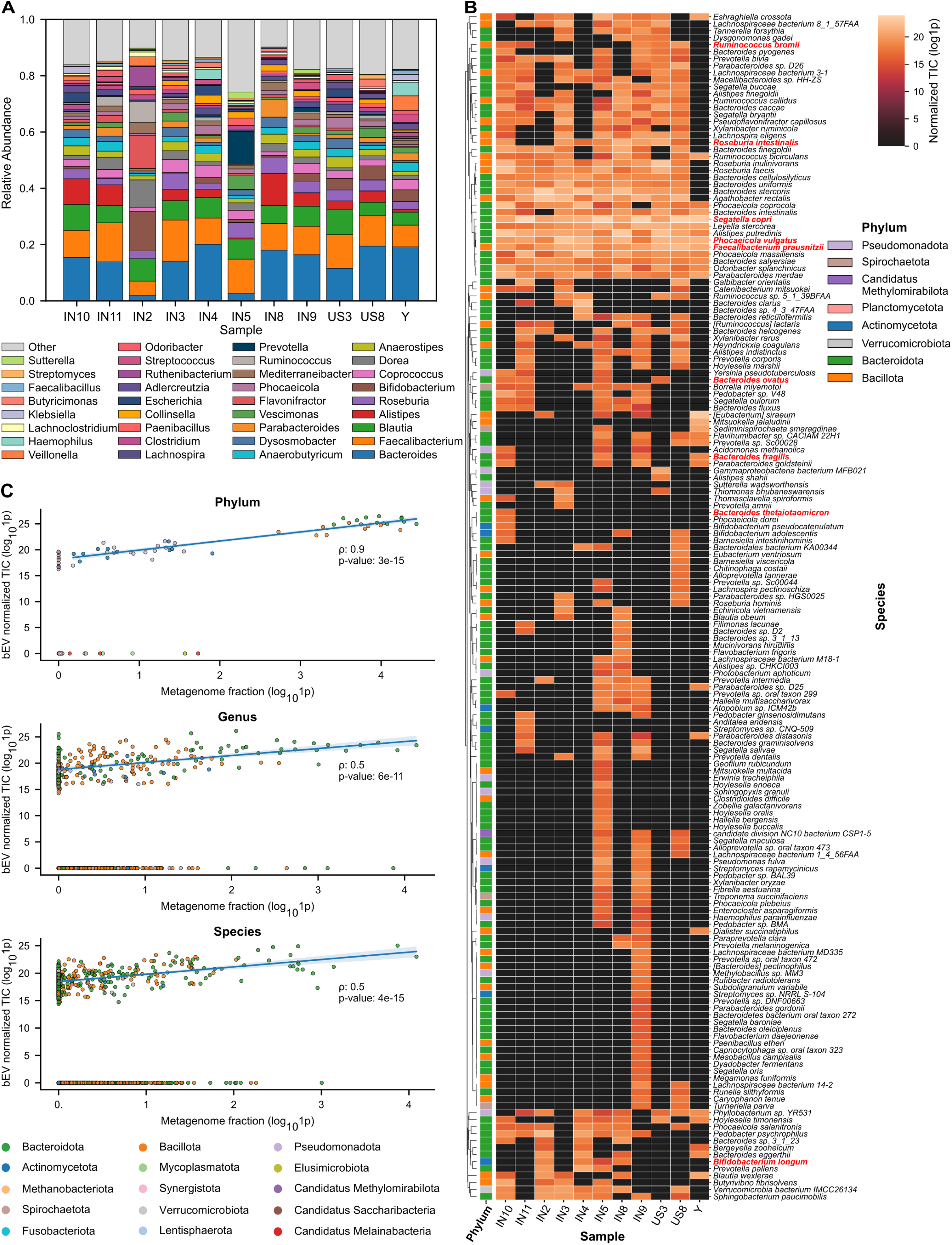
Taxonomic profiling of stool and stool-derived bEVs. (A) Relative taxonomic abundance of stool metagenomic reads using Kraken2. (B) Taxonomic distribution of bEV-associated proteomes at the species level with NCBI updated names. Colors represent the normalized total ion count (TIC) values. Color bar (left) represents the phylum per bacteria. Species names appearing in red are those from which bEVs were isolated in later experiments. (C) Spearman correlation of microbial taxa detected in the stool metagenomes and stool-derived bEV proteomes.

**Supplementary Figure 2.**
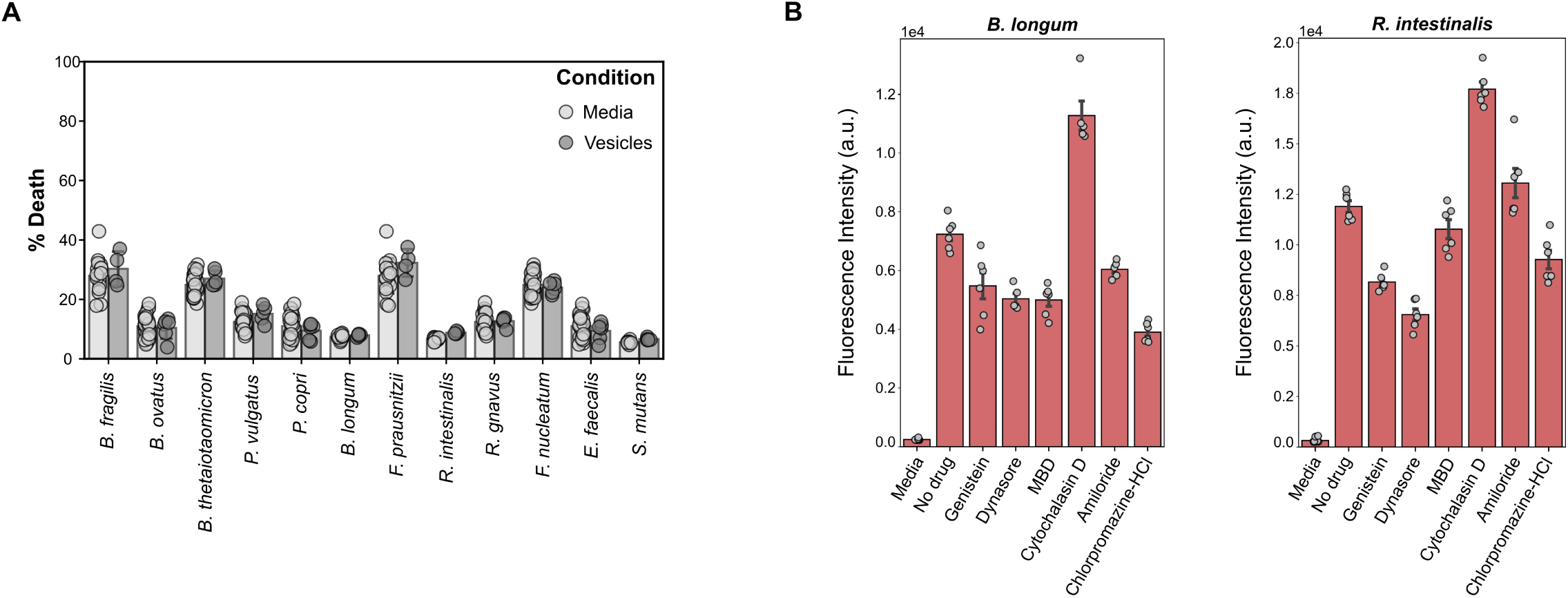
Validation controls for isolate-derived bEVs. (A) Quantification of cell death in Caco-2 cells following vesicle treatment. Cell death was assessed by flow cytometry using a Fixable Viability Dye (Invitrogen). (n=3-6) (B) Comparison of fluorescence intensity of Caco-2 cells treated with B. longum and R. intestinalis in no-drug and with drug conditions. (n=4-6, standard error of mean)

**Supplementary Figure 3.**
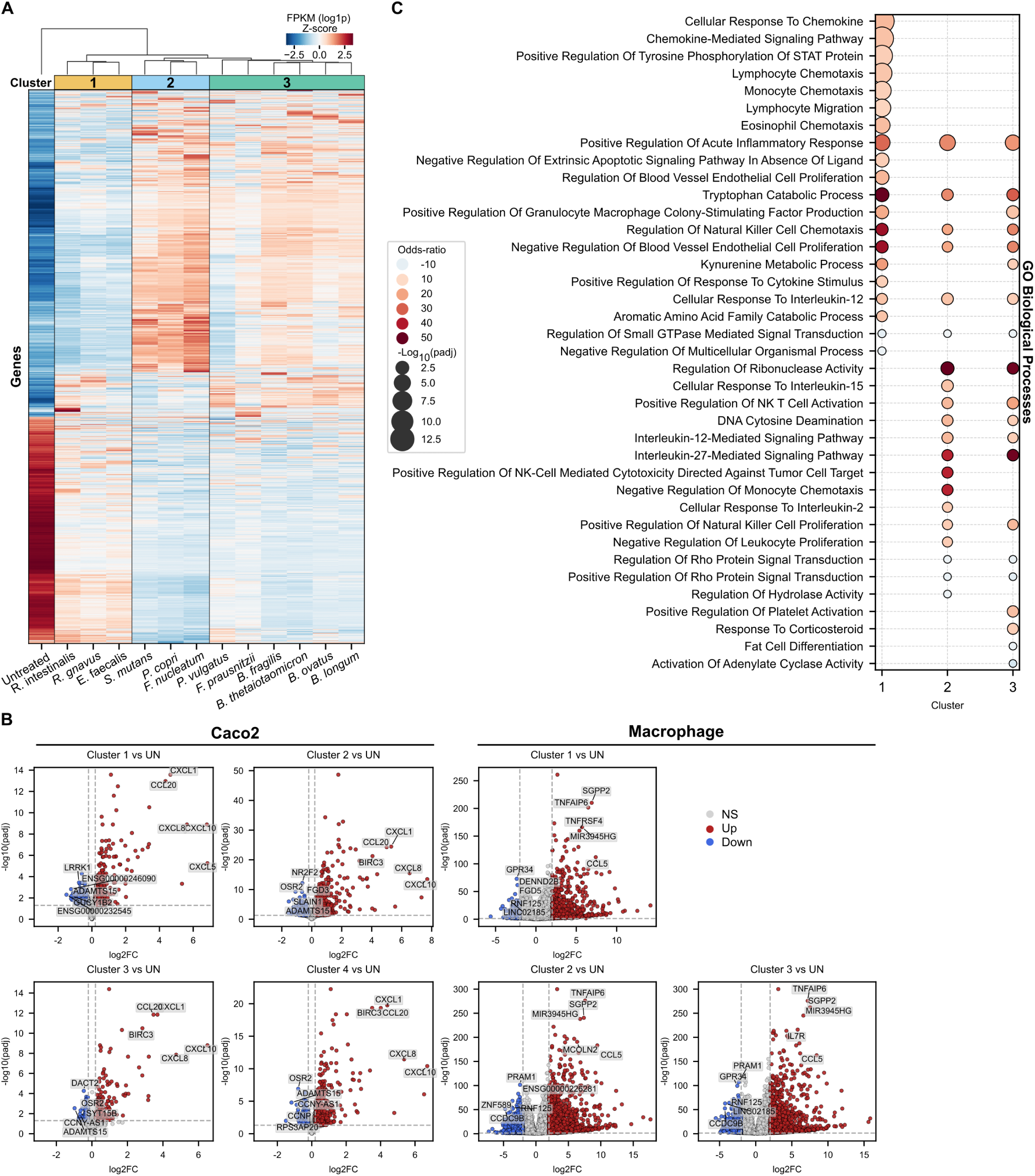
Transcriptomics analysis of epithelial cells and macrophages treated with bEVs. (A) Hierarchical clustering (clustermap) of macrophage samples following treatment with bEVs derived from distinct gut bacterial species. Color represents the FPKM-normalized z-score per gene. (B) Volcano plots showing differentially expressed genes within each identified cluster (shown in Figure 3B for Caco2 cells and Supplementary Figure 3Afor macrophages) relative to untreated controls for Caco-2 and macrophage. Annotations indicate the top five upregulated and top five downregulated genes ranked by both Iog2(fold change) and -Iog10(adjusted p-value). (C) Top 12 Gene Ontology Biological Process (GO- BP) pathways enriched in the clusters shown in (A). Color represents the odds-ratio, size represents the -Iog10(adjusted p-

**Supplementary Figure 4.**
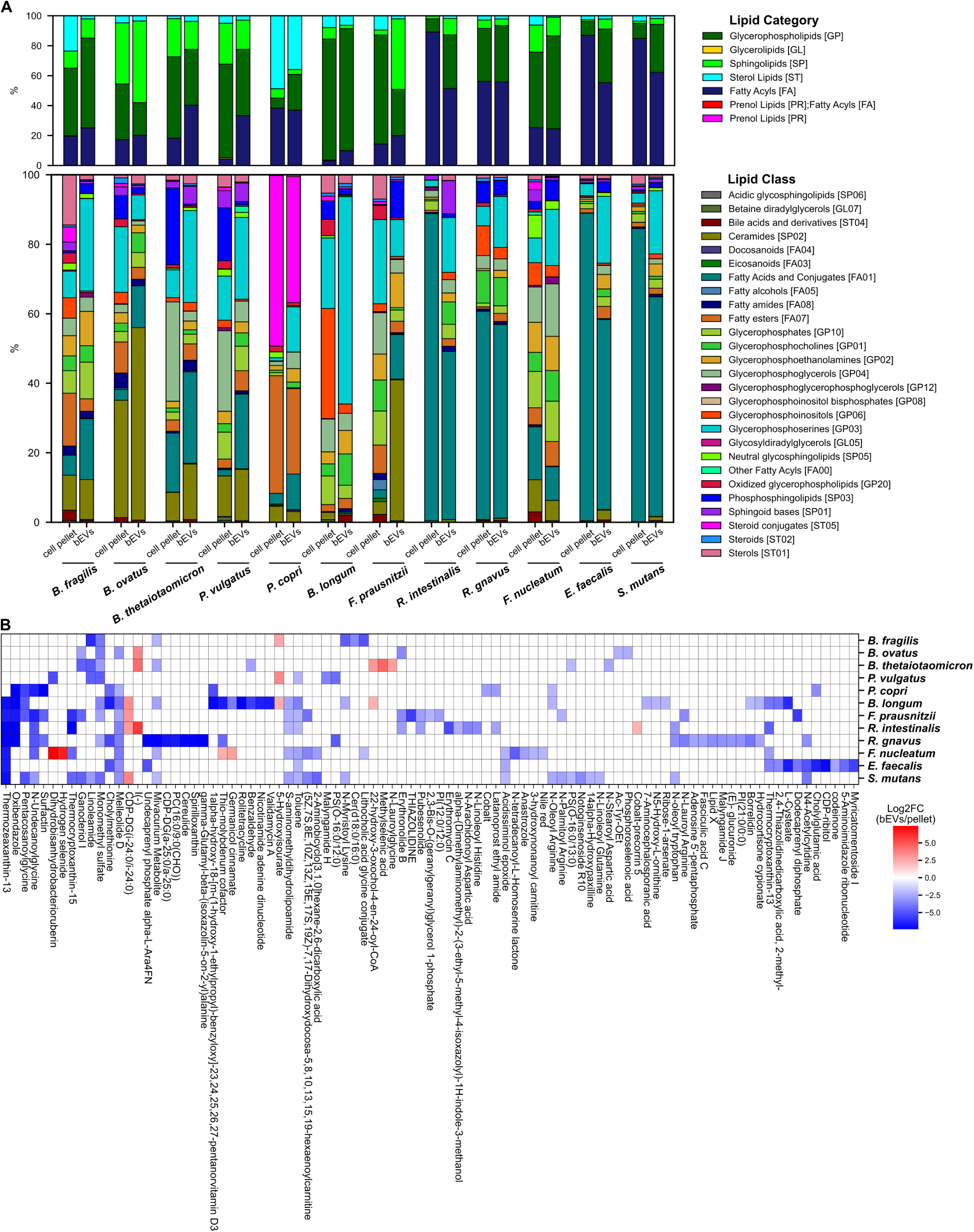
LC-MS based lipidomic and metabolomic profiling of isolated-derived bEVs. (A) Relative composition by intensity of lipid categories (upper panel) and their constituent lipid classes (lower panel) across different bEV species. (B) Top 10 most enriched and most depleted metabolites in bEVs relative to cell pellets. Color represents the Iog2(fold change).

**Supplementary Figure 5.**
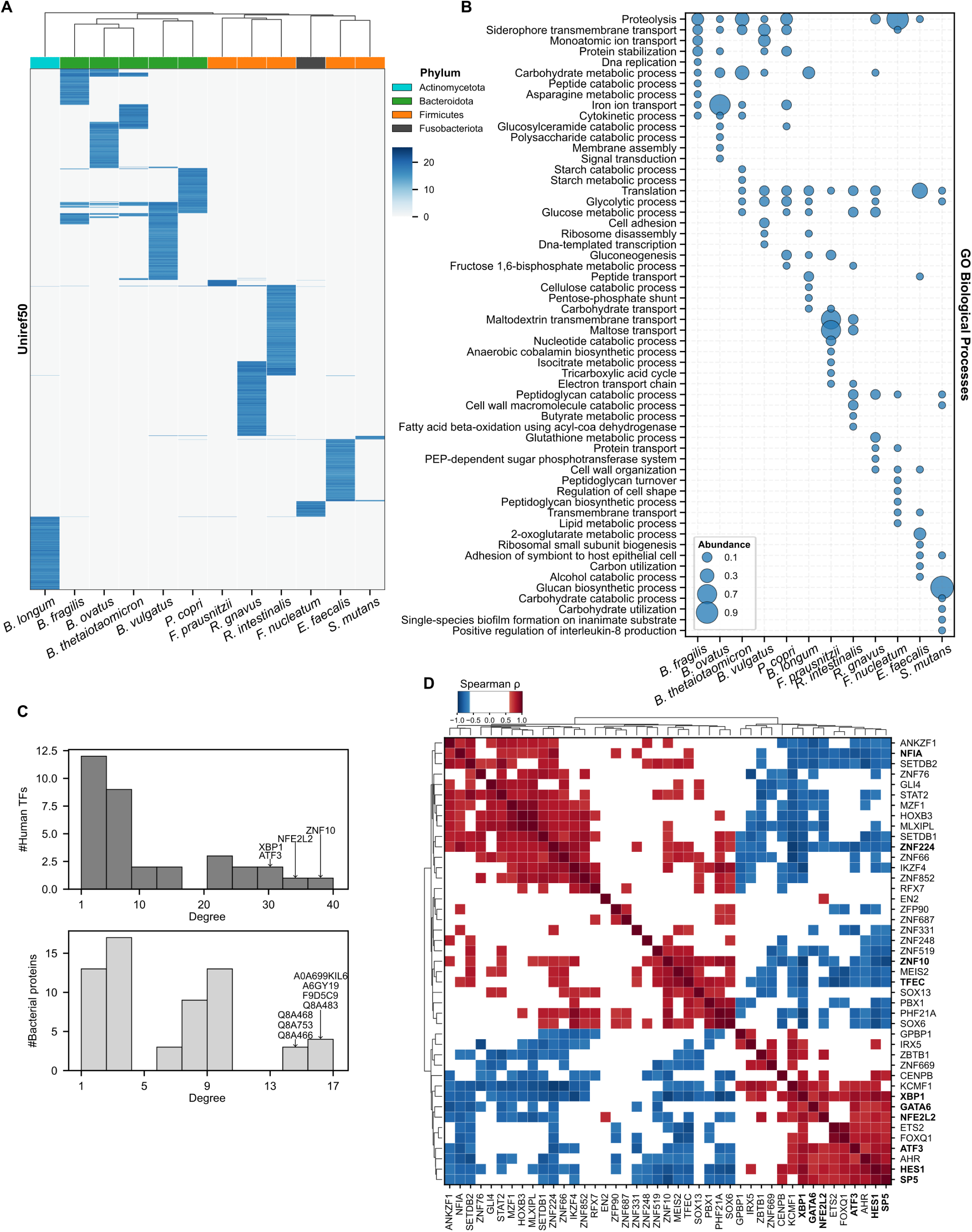
Correlation of bEV proteome cargo with the transcriptomic response of Caco-2 cells. **(A)** Hierarchical clustering of bEV protein cargo based on UniRef50 clusters. Color represents phylum. (B) Top 10 enriched Gene Ontology (GO) Biological Process terms associated with bEV cargo proteins. Size represents median abundance across replicates. (C) Degree distributions of the bacterial protein-host TF correlation network, showing the number of bacterial proteins associated with each host TF (top) and the number of host transcription factors (TFs) associated with each bacterial protein (bottom). (D) Co-expression analysis of transcription factors whose expression significantly correlated with bEV proteome cargo. Color represents the Spearman p value.

## References

1. Jones, E. J. et al. The Uptake, Trafficking, and Biodistribution of Bacteroides thetaiotaomicron Generated Outer Membrane Vesicles. Frontiers in Microbiology 11, (2020).

2. Hong, M. et al. Fusobacterium nucleatum aggravates rheumatoid arthritis through FadA-containing outer membrane vesicles. Cell Host & Microbe 31, 798–810.e7 (2023).

3. Toyofuku, M., Nomura, N. & Eberl, L. Types and origins of bacterial membrane vesicles. Nat Rev Microbiol 17, 13–24 (2019).

4. Lekmeechai, S. et al. Helicobacter pylori Outer Membrane Vesicles Protect the Pathogen From Reactive Oxygen Species of the Respiratory Burst. Front. Microbiol. 9, (2018).

5. Wai, S. N. et al. Vesicle-Mediated Export and Assembly of Pore-Forming Oligomers of the Enterobacterial ClyA Cytotoxin. Cell 115, 25–35 (2003).

6. Horstman, A. L. & Kuehn, M. J. Enterotoxigenic Escherichia coli Secretes Active Heat-labile Enterotoxin via Outer Membrane Vesicles *. Journal of Biological Chemistry 275, 12489–12496 (2000).

7. Zheng, X. et al. Fusobacterium nucleatum extracellular vesicles are enriched in colorectal cancer and facilitate bacterial adhesion. Science Advances 10, eado0016 (2024).

8. Waller, T. et al. Porphyromonas gingivalis Outer Membrane Vesicles Induce Selective Tumor Necrosis Factor Tolerance in a Toll-Like Receptor 4- and mTOR-Dependent Manner. Infection and Immunity 84, 1194–1204 (2016).

9. O’Donoghue, E. J. & Krachler, A. M. Mechanisms of outer membrane vesicle entry into host cells. Cell Microbiol 18, 1508–1517 (2016).

10. Vanaja, S. K. et al. Bacterial Outer Membrane Vesicles Mediate Cytosolic Localization of LPS and Caspase-11 Activation. Cell 165, 1106–1119 (2016).

11. Tiku, V. et al. Outer membrane vesicles containing OmpA induce mitochondrial fragmentation to promote pathogenesis of Acinetobacter baumannii. Sci Rep 11, 618 (2021).

12. Shen, Y. et al. Outer Membrane Vesicles of a Human Commensal Mediate Immune Regulation and Disease Protection. Cell Host & Microbe 12, 509–520 (2012).

13. Hickey, C. A. et al. Colitogenic *Bacteroides thetaiotaomicron* Antigens Access Host Immune Cells in a Sulfatase-Dependent Manner via Outer Membrane Vesicles. Cell Host & Microbe 17, 672–680 (2015).

14. Lázaro-Ibáñez, E. et al. Selection of Fluorescent, Bioluminescent, and Radioactive Tracers to Accurately Reflect Extracellular Vesicle Biodistribution in Vivo. ACS Nano 15, 3212–3227 (2021).

15. Edgar, R. C. Accuracy of taxonomy prediction for 16S rRNA and fungal ITS sequences. PeerJ 6, e4652 (2018).

16. Li, C.-C. et al. Characterization of markers, functional properties, and microbiome composition in human gut-derived bacterial extracellular vesicles. Gut Microbes 15, 2288200 (2023).

17. Fonseca, S. et al. Extracellular vesicles produced by the human gut commensal bacterium Bacteroides thetaiotaomicron elicit anti-inflammatory responses from innate immune cells. Front. Microbiol. 13, (2022).

18. Weinberger, V. et al. Proteomic and metabolomic profiling of extracellular vesicles produced by human gut archaea. Nat Commun 16, 5094 (2025).

19. Thapa, H. B. et al. Characterization of the Inflammatory Response Evoked by Bacterial Membrane Vesicles in Intestinal Cells Reveals an RIPK2-Dependent Activation by Enterotoxigenic Escherichia coli Vesicles. Microbiology Spectrum 11, e01115–23 (2023).

20. Schmieder, R. & Edwards, R. Quality control and preprocessing of metagenomic datasets. Bioinformatics 27, 863–864 (2011).

21. Rotmistrovsky: BMTagger: Best Match Tagger for removing… - Google Scholar. https://scholar.google.com/scholar_lookup?title=BMTagger%3A+Best+Match+Tagger+for+removing+human+reads+from+metagenomics+datasets&author=K+Rotmistrovsky&author=R+Agarwala&publication_year=2011.

22. Bolger, A. M., Lohse, M. & Usadel, B. Trimmomatic: a flexible trimmer for Illumina sequence data. Bioinformatics 30, 2114–2120 (2014).

23. Bankevich, A. et al. SPAdes: A New Genome Assembly Algorithm and Its Applications to Single-Cell Sequencing. J Comput Biol 19, 455–477 (2012).

24. Nurk, S., Meleshko, D., Korobeynikov, A. & Pevzner, P. A. metaSPAdes: a new versatile metagenomic assembler. Genome Res 27, 824–834 (2017).

25. Hyatt, D. et al. Prodigal: prokaryotic gene recognition and translation initiation site identification. BMC Bioinformatics 11, 119 (2010).

26. Buchfink, B., Xie, C. & Huson, D. H. Fast and sensitive protein alignment using DIAMOND. Nat Methods 12, 59–60 (2015).

27. Li, W. & Godzik, A. Cd-hit: a fast program for clustering and comparing large sets of protein or nucleotide sequences. Bioinformatics 22, 1658–1659 (2006).

28. Almeida, A. et al. A unified catalog of 204,938 reference genomes from the human gut microbiome. Nat Biotechnol 39, 105–114 (2021).

29. The UniProt Consortium. UniProt: the Universal Protein Knowledgebase in 2023. Nucleic Acids Res 51, D523–D531 (2023).

30. Cheng, K. et al. MetaLab: an automated pipeline for metaproteomic data analysis. Microbiome 5, 157 (2017).

31. Supek, F., Bošnjak, M., Škunca, N. & Šmuc, T. REVIGO Summarizes and Visualizes Long Lists of Gene Ontology Terms. PLOS ONE 6, e21800 (2011).

32. Ye, J. et al. Oral SMEDDS promotes lymphatic transport and mesenteric lymph nodes target of chlorogenic acid for effective T-cell antitumor immunity. J Immunother Cancer 9, (2021).

33. Cha, M. et al. Efficient Labeling of Vesicles with Lipophilic Fluorescent Dyes via the Salt-Change Method. Anal. Chem. 95, 5843–5849 (2023).

34. Gut commensal microbiota drive tailored macrophage responses: Cell Reports. https://www.cell.com/cell-reports/fulltext/S2211-1247(25)00928-3.

35. Kim, D., Paggi, J. M., Park, C., Bennett, C. & Salzberg, S. L. Graph-based genome alignment and genotyping with HISAT2 and HISAT-genotype. Nat Biotechnol 37, 907–915 (2019).

36. Li, H. et al. The Sequence Alignment/Map format and SAMtools. Bioinformatics 25, 2078–2079 (2009).

37. Liao, Y., Smyth, G. K. & Shi, W. featureCounts: an efficient general purpose program for assigning sequence reads to genomic features. Bioinformatics 30, 923–930 (2014).

38. Love, M. I., Huber, W. & Anders, S. Moderated estimation of fold change and dispersion for RNA-seq data with DESeq2. Genome Biol 15, 550 (2014).

39. Kuleshov, M. V. et al. Enrichr: a comprehensive gene set enrichment analysis web server 2016 update. Nucleic Acids Res 44, W90–W97 (2016).

40. Planck, K. A. & Rhee, K. Metabolomics of Mycobacterium tuberculosisMycobacterium tuberculosis (M. tuberculosis). in Mycobacteria Protocols (eds Parish, T. & Kumar, A.) 579–593 (Springer US, New York, NY, 2021). doi:10.1007/978-1-0716-1460-0_25.

41. Tautenhahn, R., Patti, G. J., Rinehart, D. & Siuzdak, G. XCMS Online: a web-based platform to process untargeted metabolomic data. Anal Chem 84, 5035–5039 (2012).

42. Kessner, D., Chambers, M., Burke, R., Agus, D. & Mallick, P. ProteoWizard: open source software for rapid proteomics tools development. Bioinformatics 24, 2534–2536 (2008).

43. Conroy, M. J. et al. LIPID MAPS: update to databases and tools for the lipidomics community. Nucleic Acids Res 52, D1677–D1682 (2024).

44. Gil-de-la-Fuente, A. et al. CEU Mass Mediator 3.0: A Metabolite Annotation Tool. J. Proteome Res. 18, 797–802 (2019).

45. Hughes, C. S. et al. Single-pot, solid-phase-enhanced sample preparation for proteomics experiments. Nat Protoc 14, 68–85 (2019).

46. Frankenfield, A. M., Ni, J., Ahmed, M. & Hao, L. Protein Contaminants Matter: Building Universal Protein Contaminant Libraries for DDA and DIA Proteomics. J. Proteome Res. 21, 2104–2113 (2022).

47. Label-Free and Standard-Free Absolute Quantitative Proteomics Using the “Total Protein” and “Proteomic Ruler” Approaches. in Methods in Enzymology vol. 585 49–60 (Academic Press, 2017).

48. Brown, S. T. et al. Bridges-2: A Platform for Rapidly-Evolving and Data Intensive Research. in Practice and Experience in Advanced Research Computing 2021: Evolution Across All Dimensions 1–4 (Association for Computing Machinery, New York, NY, USA, 2021). doi:10.1145/3437359.3465593.

49. Accurate structure prediction of biomolecular interactions with AlphaFold 3 | Nature. https://www.nature.com/articles/s41586-024-07487-w.

50. Fast and accurate protein structure search with Foldseek | Nature Biotechnology. https://www.nature.com/articles/s41587-023-01773-0.

51. Lambert, S. A. et al. The Human Transcription Factors. Cell 172, 650–665 (2018).

52. Müller-Dott, S. et al. Expanding the coverage of regulons from high-confidence prior knowledge for accurate estimation of transcription factor activities. Nucleic Acids Res 51, 10934–10949 (2023).

53. Garcia-Alonso, L., Holland, C. H., Ibrahim, M. M., Turei, D. & Saez-Rodriguez, J. Benchmark and integration of resources for the estimation of human transcription factor activities. Genome Res. 29, 1363–1375 (2019).

54. Badia-i-Mompel, P. et al. decoupleR: ensemble of computational methods to infer biological activities from omics data. Bioinformatics Advances 2, vbac016 (2022).

55. Yu, G. & He, Q.-Y. ReactomePA: an R/Bioconductor package for reactome pathway analysis and visualization. Mol. BioSyst. 12, 477–479 (2016).

56. Coelho, C. & Casadevall, A. Answers to naysayers regarding microbial extracellular vesicles. Biochem Soc Trans 47, 1005–1012 (2019).

57. McMillan, H. M. & Kuehn, M. J. The extracellular vesicle generation paradox: a bacterial point of view. EMBO J 40, EMBJ2021108174 (2021).

58. Quévrain, E. et al. Identification of an anti-inflammatory protein from Faecalibacterium prausnitzii, a commensal bacterium deficient in Crohn’s disease. 10.1136/gutjnl-2014-307649 (2016) doi:10.1136/gutjnl-2014-307649.

59. Nakjang, S., Ndeh, D. A., Wipat, A., Bolam, D. N. & Hirt, R. P. A Novel Extracellular Metallopeptidase Domain Shared by Animal Host-Associated Mutualistic and Pathogenic Microbes. PLOS ONE 7, e30287 (2012).

60. Ding, X. et al. Bacteroides fragilis promotes chemoresistance in colorectal cancer, and its elimination by phage VA7 restores chemosensitivity. Cell Host & Microbe 33, 941–956.e10 (2025).

61. Wang, Y., Ngo, V. L., Zou, J. & Gewirtz, A. T. Commensal bacterial outer membrane protein A induces interleukin-22 production. Cell Reports 43, (2024).

62. Weinberger, V. et al. Proteomic and metabolomic profiling of extracellular vesicles produced by human gut archaea. Nat Commun 16, 5094 (2025).

63. Chelakkot, C. et al. Akkermansia muciniphila-derived extracellular vesicles influence gut permeability through the regulation of tight junctions. Exp Mol Med 50, e450–e450 (2018).

64. Park, H. S. et al. Clinical features and KRAS mutation in colorectal cancer with bone metastasis. Sci Rep 10, 21180 (2020).

65. Nam, J. H. et al. Osteoporosis Is Associated with an Increased Risk of Colorectal Adenoma and High-Risk Adenoma: A Retrospective, Multicenter, Cross-Sectional, Case-Control Study. Gut and Liver 16, 269–276 (2022).

66. MacNair, C. R. et al. Outer Membrane Vesicles Hijack TIM-1 for Cellular Uptake. 2025.12.22.696083 Preprint at 10.64898/2025.12.22.696083 (2025).

67. Preet, R. et al. Gut commensal Bifidobacterium-derived extracellular vesicles modulate the therapeutic effects of anti-PD-1 in lung cancer. Nat Commun 16, 3500 (2025).

68. He, L., Sayers, E. J., Watson, P. & Jones, A. T. Contrasting roles for actin in the cellular uptake of cell penetrating peptide conjugates. Sci Rep 8, 7318 (2018).

69. Fu, X. et al. Precise design strategies of nanomedicine for improving cancer therapeutic efficacy using subcellular targeting. Sig Transduct Target Ther 5, 262 (2020).

70. Hegarty, L. M., Jones, G.-R. & Bain, C. C. Macrophages in intestinal homeostasis and inflammatory bowel disease. Nat Rev Gastroenterol Hepatol 20, 538–553 (2023).

71. Xiong, Y. et al. Bacteroides Fragilis Transplantation Reverses Reproductive Senescence by Transporting Extracellular Vesicles Through the Gut-Ovary Axis. 10.1002/advs.202409740 doi:10.1002/advs.202409740.

72. Nie, X. et al. Bifidobacterium longum NSP001-derived extracellular vesicles ameliorate ulcerative colitis by modulating T cell responses in gut microbiota-(in)dependent manners. npj Biofilms Microbiomes 11, 27 (2025).

73. Moosavi, S. M., Akhavan Sepahi, A., Mousavi, S. F., Vaziri, F. & Siadat, S. D. The effect of Faecalibacterium prausnitzii and its extracellular vesicles on the permeability of intestinal epithelial cells and expression of PPARs and ANGPTL4 in the Caco-2 cell culture model. J Diabetes Metab Disord 19, 1061–1069 (2020).

74. Krämer, A., Green, J., Pollard, J., Jr & Tugendreich, S. Causal analysis approaches in Ingenuity Pathway Analysis. Bioinformatics 30, 523–530 (2014).

75. Holt, S. E. et al. Functional requirement of p23 and Hsp90 in telomerase complexes. Genes Dev 13, 817–826 (1999).

76. Zhu, X. et al. Downregulated KLF4, induced by m6A modification, aggravates intestinal barrier dysfunction in inflammatory bowel disease. Cell. Mol. Life Sci. 81, 470 (2024).

77. Fevr, T., Robine, S., Louvard, D. & Huelsken, J. Wnt/β-Catenin Is Essential for Intestinal Homeostasis and Maintenance of Intestinal Stem Cells. Mol Cell Biol 27, 7551–7559 (2007).

78. Mauer, J. et al. Interleukin-6 signaling promotes alternative macrophage activation to limit obesity-associated insulin resistance and endotoxemia. Nat Immunol 15, 423–430 (2014).

79. Groß, R. et al. Phosphatidylserine-exposing extracellular vesicles in body fluids are an innate defence against apoptotic mimicry viral pathogens. Nat Microbiol 9, 905–921 (2024).

80. Xu, Q. et al. A Distinct Type of Pilus from the Human Microbiome. Cell 165, 690–703 (2016).

81. Spiga, L. et al. Iron acquisition by a commensal bacterium modifies host nutritional immunity during *Salmonella* infection. Cell Host & Microbe 31, 1639–1654.e10 (2023).

82. Harvey, K. L., Jarocki, V. M., Charles, I. G. & Djordjevic, S. P. The Diverse Functional Roles of Elongation Factor Tu (EF-Tu) in Microbial Pathogenesis. Front Microbiol 10, 2351 (2019).

83. Granato, D. et al. Cell Surface-Associated Elongation Factor Tu Mediates the Attachment of Lactobacillus johnsonii NCC533 (La1) to Human Intestinal Cells and Mucins. Infection and Immunity 72, 2160–2169 (2004).

84. Cui, G., Li, P., Wu, R. & Lin, H. Streptococcus mutans membrane vesicles inhibit the biofilm formation of Streptococcus gordonii and Streptococcus sanguinis. AMB Expr 12, 154 (2022).

85. Briaud, P. et al. Temperature Influences the Composition and Cytotoxicity of Extracellular Vesicles in Staphylococcus aureus. mSphere 6, e00676–21.

86. Macdonald, I. A. & Kuehn, M. J. Stress-induced outer membrane vesicle production by Pseudomonas aeruginosa. https://hdl.handle.net/10161/10655 (2013).

87. Bos, J., Cisneros, L. H. & Mazel, D. Real-time tracking of bacterial membrane vesicles reveals enhanced membrane traffic upon antibiotic exposure. Science Advances 7, eabd1033 (2021).

88. Toussi, D. N., Liu, X. & Massari, P. The FomA Porin from Fusobacterium nucleatum Is a Toll-Like Receptor 2 Agonist with Immune Adjuvant Activity. Clinical and Vaccine Immunology 19, 1093–1101 (2012).

89. Sartorio, M. G., Pardue, E. J., Scott, N. E. & Feldman, M. F. Human gut bacteria tailor extracellular vesicle cargo for the breakdown of diet- and host-derived glycans. Proceedings of the National Academy of Sciences 120, e2306314120 (2023).

90. Chu, H. et al. Gene-microbiota interactions contribute to the pathogenesis of inflammatory bowel disease. Science 352, 1116–1120 (2016).

91. Ye, L. et al. F. prausnitzii-derived extracellular vesicles attenuate experimental colitis by regulating intestinal homeostasis in mice. Microb Cell Fact 22, 235 (2023).

92. Zhang, X. et al. Metaproteomics reveals associations between microbiome and intestinal extracellular vesicle proteins in pediatric inflammatory bowel disease. Nat Commun 9, 2873 (2018).

93. Erttmann, S. F. et al. The gut microbiota prime systemic antiviral immunity via the cGAS-STING-IFN-I axis. Immunity 55, 847–861.e10 (2022).

